# Nonreciprocal and Conditional Cooperativity Directs the Pioneer Activity of Pluripotency Transcription Factors

**DOI:** 10.1101/633826

**Authors:** Sai Li, Eric Bo Zheng, Li Zhao, Shixin Liu

**Affiliations:** Laboratory of Nanoscale Biophysics and Biochemistry, The Rockefeller University, New York, NY 10065, USA; Laboratory of Evolutionary Genetics and Genomics, The Rockefeller University, New York, NY 10065, USA

## Abstract

Cooperative binding of transcription factors (TFs) to chromatin orchestrates gene expression programming and cell fate specification. However the biophysical principles of TF cooperativity remain incompletely understood. Here we use single-molecule fluorescence microscopy to study the partnership between Sox2 and Oct4, two core members of the pluripotency gene regulatory network. We find that Sox2’s pioneer activity (the ability to target DNA inside nucleosomes) is strongly affected by the translational and rotational positioning of its binding motif, while Oct4 can access nucleosomal sites with equal capacities. Furthermore, the Sox2-Oct4 pair displays nonreciprocal cooperativity, with Oct4 modulating Sox2’s binding to the nucleosome but not vice versa. Such cooperativity is conditional upon the composite motif residing at specific nucleosomal locations. These results reveal that pioneer factors possess distinct properties of nucleosome targeting and suggest that the same set of TFs may differentially regulate transcriptional activity in a gene-specific manner on the basis of their motif positioning in the nucleosomal context.

## INTRODUCTION

Transcription factors (TFs) access and interpret the genome by recognizing specific DNA sequences and regulating the transcriptional activity of selected sets of genes (Lambert et al., 2018; Ptashne and Gann, 2002). In eukaryotic nuclei, genomic DNA is wrapped around histone octamers, forming nucleosome building blocks and higher-order chromatin structures (Luger et al., 2012; McGinty and Tan, 2015). The compaction of DNA into chromatin often occludes TFs from their cognate binding motifs, thus constituting an important regulatory layer of gene expression control and cell identity determination (Li et al., 2007; Segal and Widom, 2009). A subset of TFs, known as pioneer factors (PFs), possesses the ability to access nucleosomal DNA and closed chromatin, which further recruits other chromatin-binding proteins and transcription machinery to their target sites and initiate transcriptional reprogramming and cell fate transitions (Zaret and Mango, 2016).

TF pairs can exhibit cooperative binding behavior, i.e., binding of one factor to DNA facilitates targeting of the other (Morgunova and Taipale, 2017). Such cooperativity can be mediated by direct TF-TF interaction or through the DNA substrate (Jolma et al., 2015; Kim et al., 2013). Alternatively, due to the competition between TFs and nucleosomes for DNA binding, TF-TF cooperativity can be manifested indirectly and, often nonspecifically, in the context of chromatin (Adams and Workman, 1995; Mirny, 2010; Polach and Widom, 1996; Vashee et al., 1998). In this scenario, PFs are the first to engage and open up closed chromatin, making it more accessible to other TFs (Sartorelli and Puri, 2018; Zaret and Carroll, 2011).

One of the most prominent examples of TFs shaping gene expression pattern is the “Yamanaka” factors (Sox2, Oct4, Klf4, and c-Myc), which can convert mammalian somatic cells into induced pluripotent stem cells (Takahashi and Yamanaka, 2006). It was shown that Sox2, Oct4, and Klf4, but not c-Myc, can function as PFs by binding to nucleosomes *in vitro* and silent, DNase-resistant chromatin *in vivo* (Soufi et al., 2012; Soufi et al., 2015). During reprogramming towards pluripotency, these factors cooperatively target selected enhancers and activate or repress the expression of distinct genes in a stage-specific manner (Chronis et al., 2017; Soufi et al., 2012).

Among these reprogramming factors, Sox2 and Oct4 are also core members of the transcriptional regulatory network that governs embryogenesis and the maintenance of embryonic stem cells (Li and Belmonte, 2017; Rizzino and Wuebben, 2016). Sox2 contains an HMG domain that binds to the minor groove of the DNA helix, whereas Oct4 harbors a bipartite POU domain that interacts with the DNA major groove, allowing the formation of Sox2:Oct4:DNA ternary complexes with the TF pair binding to adjacent motifs (Remenyi et al., 2003; Williams et al., 2004). ChIP-seq experiments revealed that Sox2 and Oct4 co-occupy the cis-regulatory elements of a large number of target genes (Boyer et al., 2005; Chen et al., 2008; Whyte et al., 2013), suggesting that this TF pair works synergistically to regulate gene expression. Indeed, juxtaposed HMG:POU composite motifs are found upstream of many pluripotency-associated genes (Ambrosetti et al., 1997; Nishimoto et al., 1999; Okumura-Nakanishi et al., 2005; Rodda et al., 2005; Tomioka et al., 2002).

Despite extensive research, the capacity of and mutual relationship between Sox2 and Oct4, and PFs in general, in targeting nucleosomal DNA remains a matter of debate (Chronis et al., 2017; Franco et al., 2015; Iwafuchi-Doi et al., 2016; Soufi et al., 2015; Swinstead et al., 2016). Recent data suggested that the pioneer activity of a given TF may be conditional and dependent on the local chromatin environment (King and Klose, 2017; Liu and Kraus, 2017; Soufi et al., 2015; Swinstead et al., 2016). For example, it was shown that the binding of Sox2 to a subset of genomic loci is dependent on another chromatin-binding protein PARP-1, and that such dependence is correlated to the rotational positioning of the Sox2 motif in the nucleosome (Liu and Kraus, 2017). Popular methods of choice for studying TF-chromatin interaction such as genome-wide binding and bulk biochemical assays lack sufficient temporal resolution to inform the time order of binding events (usually occurring on the order of seconds) by multiple TFs. Single-particle tracking experiments in living cells can capture the dynamic nature of TF binding (Chen et al., 2014; Lam et al., 2012; White et al., 2016), but the DNA sequence and chromatin state of the binding targets in these assays are usually not well defined. By contrast, *in vitro* single-molecule measurements allow for precise control of the substrates and have been employed to provide quantitative information on the binding/dissociation kinetics of TFs and chromatin regulators (Choi et al., 2017; Gibson et al., 2017; Kilic et al., 2015; Luo et al., 2014).

In this work, we used single-molecule fluorescence microscopy to measure the binding dynamics of Sox2 and Oct4—both individually and in combination—on a variety of DNA and nucleosome substrates. We found that, although both classified as PFs, Sox2 and Oct4 exhibit markedly distinct properties of nucleosome targeting. Sox2 strongly favors nucleosome-dyad-positioned sites over end-positioned ones, whereas Oct4 indiscriminately binds to targets at all positions. Oct4 and Sox2 are hierarchically recruited to composite nucleosomal motifs, with Oct4 predominantly engaging first. We further demonstrated that the Sox2-Oct4 pair displays nonreciprocal cooperativity, and that such cooperativity is dependent on the positioning of the composite motif with respect to the nucleosome. Consistent with these *in vitro* results, our analyses of the genomic data showed that the DNA binding sites of Sox2, but not Oct4, are preferentially located toward the center of the nucleosome. These results help to clarify the biophysical rules governing the Sox2-Oct4 partnership and suggest that genes may be differentially regulated by the same set of TFs on the basis of their motif positioning in the nucleosomal context.

## RESULTS

### Single-Molecule Analysis of Sox2 Binding to DNA

We labeled the purified full-length human Sox2 with a Cy5 fluorophore near its C-terminus (Figure S1). We then constructed a DNA template containing a 147-basepair(bp)-long 601 nucleosome positioning sequence (NPS) with a canonical Sox2 binding motif (CTTTGTT) located at its end (nucleotide #1-7), which was termed as DNA_S-End_ (Figure 1A). Cy3-labeled DNA templates were immobilized on a glass coverslip and their locations visualized by total-internal-reflection fluorescence (TIRF) microscopy (Figure 1B). Cy5-Sox2 was injected into the flow chamber at a concentration of ~ 2 nM. Binding and dissociation of individual Sox2 molecules at the DNA loci were monitored in real time. Single-molecule fluorescence trajectories allowed us to measure the residence time (*t*_bound_) of Sox2 on DNA (Figure 1C). The cumulative distribution of *t*_bound_ built from many binding events can be well-fit by a single-exponential function (Figure 1D). After correcting for dye photobleaching (Figure S2), we determined the characteristic lifetime of Sox2 binding (*τ*)—governed by the corresponding dissociation rate constant—to be 19.7 ± 2.3 s (Table S1).

**Figure 1.**
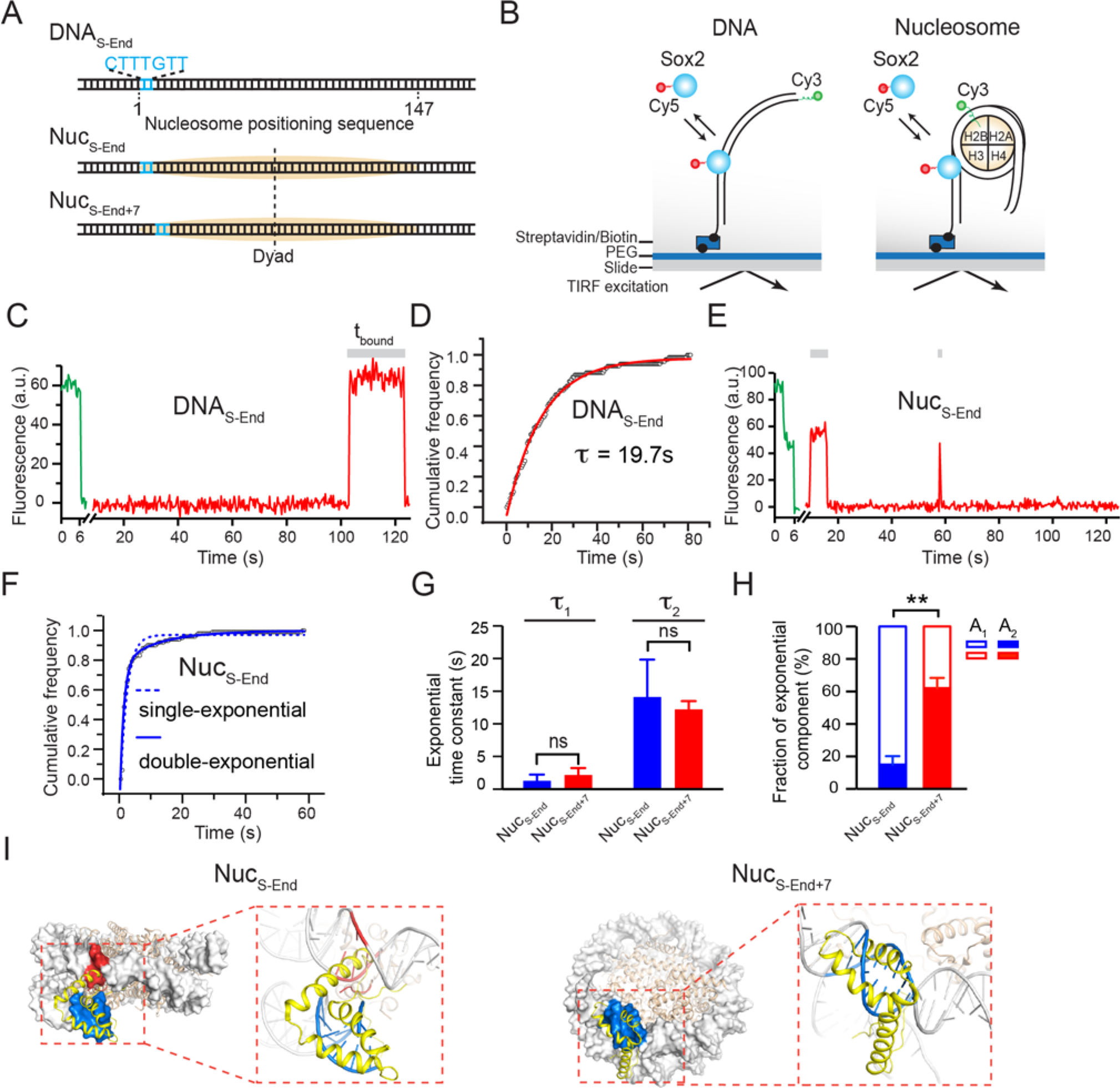
Sox2 Displays Differential Binding Kinetics on DNA and Nucleosome Substrates. (**A**) Diagrams of DNA and nucleosome substrates containing a Sox2 binding motif (blue) located near the end of a 601 nucleosome positioning sequence (orange). (**B**) Schematic of the single-molecule TF binding assay using a total-internal-reflection fluorescence microscope. (**C**) A representative fluorescence-time trajectory showing Cy5-labeled Sox2 binding to a Cy3-labeled DNA_S-End_ substrate. A 532-nm laser was first turned on briefly to locate the surface-immobilized substrates. Then a 640-nm laser was switched on to monitor Sox2 binding and dissociation. (**D**) Cumulative distribution (open circles) of the Sox2 residence time on DNA_S-End_ [*t*_bound_ as shown in (C)] and its fit to a single-exponential function y(t) = A × exp(−t/*τ*) + y_0_ (red curve). (**E**) A representative fluorescence-time trajectory showing Cy5-labeled Sox2 binding to a Cy3-labeled Nuc_S-End_ substrate. The two photobleaching steps under 532-nm excitation confirm the existence of two Cy3-labeled H2B, suggesting an intact nucleosome. (**F**) Cumulative distribution (filled circles) of the Sox2 residence time on Nuc_S-End_ and its fit to a double-exponential function y(t) = A_1_ × exp(-t/*τ*_1_) + A_2_ × exp(−t/*τ*_2_) + y_0_ (solid blue curve). The dashed blue curve displays a poor fit to a single-exponential function. (**G**) Time constants for the two exponential components (*τ*_1_ and *τ*_2_) from the double-exponential fit shown in F (blue bars for Nuc_S-End_; red bars for Nuc_S-End+7_). (**H**) Relative weights of the fast (A_1_) and slow (A_2_) exponential components for Nuc_S-End_ (blue) and Nuc_S-End+7_ (red). (**I**) The Sox2_HMG_:DNA structure (PDB: 1GT0; yellow) superimposed on the nucleosome structure (PDB: 3LZ0; grey) aligned by the DNA motif (blue), which spans nucleotide #1-7 for Nuc_S-End_ (left) and #8-14 for Nuc_S-End+7_ (right). Steric clash between Sox2 and the nucleosome is highlighted in red. Data are represented as mean ± SD.

### Sox2 Fails to Stably Bind to End-Positioned Nucleosomal DNA

Sox2 is considered as a PF, capable of targeting binding sites within nucleosome-occupied genomic regions (Zaret and Mango, 2016). To directly observe the behavior of Sox2 on nucleosomal DNA, we created a mononucleosome substrate, Nuc_S-End_, which was assembled with DNA_S-End_ and Cy3-labeled histone H2B as well as unlabeled H2A, H3, and H4 (Figure 1A). Fluorescence signal from Cy3-H2B was used to locate individual nucleosome substrates on the surface (Figure 1E). We used DNase I digestion followed by sequencing to show that Nuc_S-End_ shares an identical digestion pattern with nucleosomes assembled with the unmodified 601 NPS (Figures S3A-S3B), suggesting that nucleosome positioning is not perturbed by the engineered Sox2 binding motif.

The residence time distribution for Sox2 on Nuc_S-End_ cannot be described by a single-exponential model, but is well-fit by two exponential components: a predominant, shorter-lived population (*τ*_1_ = 1.3 s; fraction A_1_ = 85%) and a rare, longer-lived one (*τ*_2_ = 14.1 s; A_2_ = 15%) (Figures 1F-1H; Table S1). The fast component likely corresponds to the nonspecific sampling of Sox2 on chromatin substrates as reported before (Chen et al., 2014). The slow component, on the other hand, likely represents the specific interaction between Sox2 and its cognate DNA sequence. Notably, the specific binding events only constitute a minor fraction of all binding events and feature a shorter lifetime than those on bare DNA (14.1 s vs. 19.7 s), indicating that nucleosome packing makes the DNA motif less accessible to Sox2. Indeed, when we superimposed the Sox2-DNA structure with the nucleosome structure, we observed a significant steric clash of the Sox2 HMG domain against the histone H3 and the neighboring DNA gyre (Figure 1I, Nuc_S-End_). Thus, single-molecule measurements led to the unexpected finding that nucleosome wrapping can significantly affect DNA targeting by Sox2.

Next we moved the Sox2 binding motif inward (toward the nucleosome dyad) by 7 bp and created a new nucleosome substrate Nuc_S-End+7_. This change shifts the phasing of the minor groove face of the DNA motif—which is recognized by Sox2—relative to the histone octamer, such that the steric hindrance between Sox2 and the nucleosome is much reduced (Figure 1I, Nuc_S-End+7_). Accordingly, we observed a significantly larger fraction of long-lived binding events compared to Nuc_S-End_ (A_2_ = 62% for Nuc_S-End+7_ vs. 15% for Nuc_S-End_; Figure 1H), supporting the notion that these events correspond to the specific Sox2-nucleosome interaction mode.

### Sox2 Stably Engages with Binding Sites near the Nucleosome Dyad

A recent study showed that the Sox family TFs exhibit preferred binding around the nucleosome dyad region (Zhu et al., 2018). To dissect the single-molecule binding kinetics of Sox2 at the dyad, we placed the Sox2 binding motif at the center of the 601 NPS (nucleotide #72-78) and assembled a nucleosome substrate Nuc_S-Dyad_ (Figure 2A). Again, such sequence engineering did not change the overall nucleosomal organization according to the DNase digestion pattern (Figure S3C). In stark contrast to its behavior on Nuc_S-End_, Sox2 exhibited more prolonged binding to Nuc_S-Dyad_ (an average residence time of 22.2 s for Nuc_S-Dyad_ vs. 4.6 s for Nuc_S-End_). The residence time distribution for Sox2 on Nuc_S-Dyad_ is also characterized by a double-exponential function (*τ*_1_ = 2.1 s; *τ*_2_ = 36.8 s). Importantly, specific Sox2 binding events on Nuc_S-Dyad_ are both longer-lived (*τ*_2_) and more prevalent (A_2_) than those on Nuc_S-End_ (Figures 2B-2C, compare End and Dyad columns), consistent with the idea that the dyad position presents an optimal environment for Sox2 engagement. This is further supported by structural superposition that shows minimal interference imposed by histones and nucleosomal DNA on Sox2 (Figure 2D).

**Figure 2.**
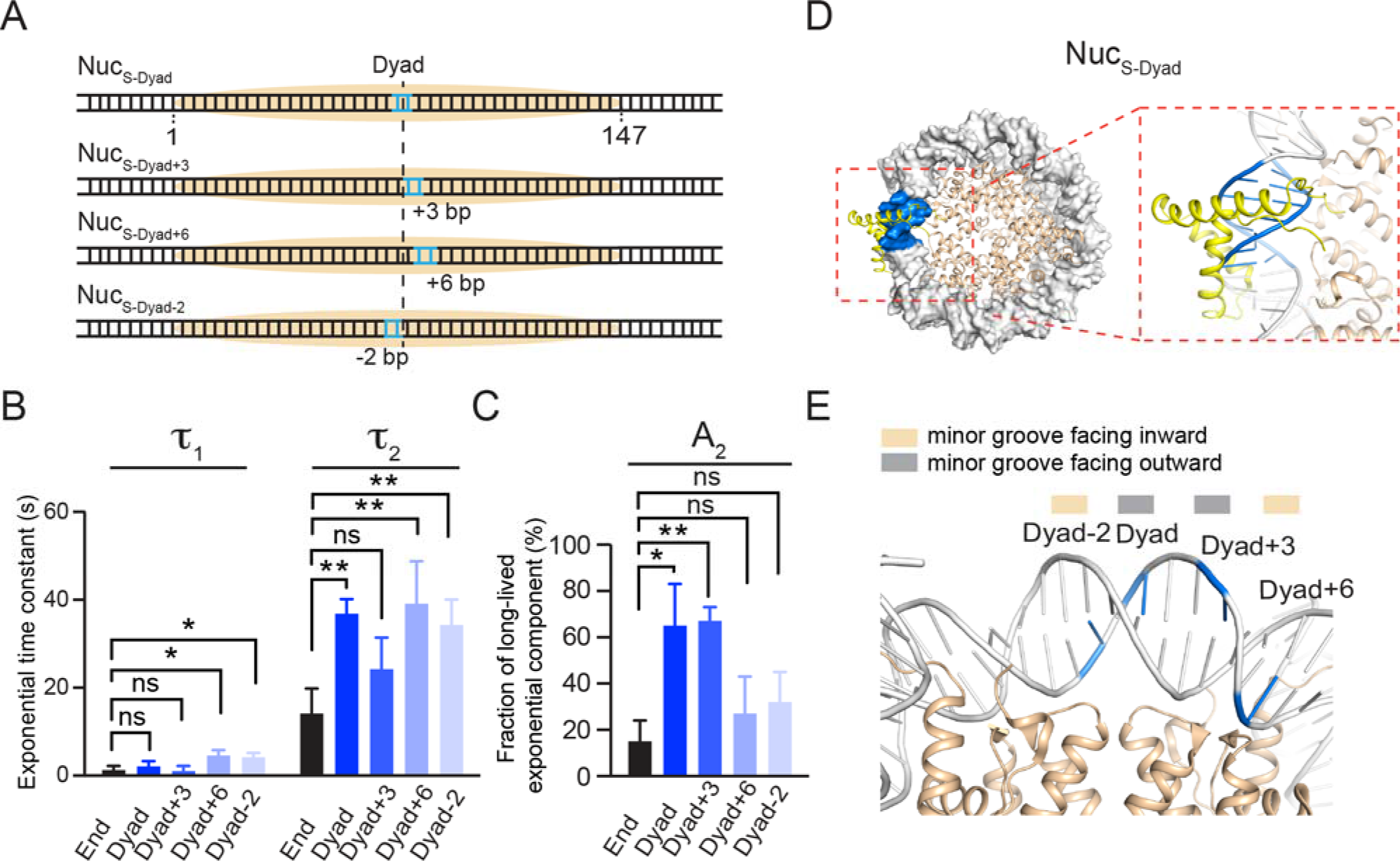
Sox2’s Pioneer Activity Near the Nucleosome Dyad Is Regulated by the Rotational Phasing of the DNA Motif. (**A**) Diagrams of nucleosome substrates harboring a Sox2 binding motif around the nucleosome dyad axis. (**B**) Time constants for the fast and slow exponential components (*τ*_1_ and *τ*_2_) that describe Sox2’s residence time on different nucleosome substrates. (**C**) Fractions of long-lived, specific Sox2 binding events (A_2_) for different nucleosome substrates. (**D**) Structural superposition illustrating the putative binding configuration of Sox2 on the Nuc_S-Dyad_ substrate. The Sox2 HMG domain and the DNA motif are shown in yellow and blue, respectively. (**E**) Zoomed-in view of the nucleosome dyad region displaying the orientation of the DNA minor groove. The midpoints of the Sox2 binding motif placed at different positions (Dyad-2, Dyad, Dyad+3, and Dyad+6) are indicated in blue. Data are represented as mean ± SD.

To evaluate whether the dyad region is in general favored by Sox2, we generated three more nucleosome substrates by shifting the Sox2 motif away from the dyad axis, either to the right by 3 bp or 6 bp, or to the left by 2 bp (Nuc_S-Dyad+3_, Nuc_S-Dyad+6_, Nuc_S-Dyad-2_, respectively; Figures 2A). We found that all dyad-positioned sites can accommodate extended Sox2 binding compared to the end-positioned site, but to different degrees (Figures 2B-2C). Nuc_S-Dyad_ and Nuc_S-Dyad+3_, in which the minor groove of the Sox2 binding motif is mostly outward facing, have the highest fractions of long-lived binding. On the other hand, the minor groove is mostly inward facing for Nuc_S-Dyad+6_ and Nuc_S-Dyad-2_, which also feature lower fractions of specific Sox2 interaction (Figures 2D-2E, S4A-S4C). Thus, even though Sox2 generally prefers the dyad region, the likelihood and lifetime of its specific nucleosome-binding mode are nonetheless influenced by the rotational phasing of the DNA motif.

The results above collectively demonstrate that Sox2’s pioneer activity is sensitively modulated by the translational and rotational positioning of its cognate DNA motif within the nucleosome.

### Oct4 Binds Equally to End- and Dyad-Positioned Nucleosomal DNA Motifs

Next we examined the behavior of the other core pluripotency TF, Oct4, on DNA and nucleosome substrates. We purified and fluorescently labeled full-length human Oct4 with Cy5 (Figure S1), incorporated an 8-bp-long Oct4 binding motif into the 601 NPS at either the end or the dyad position (DNA_O-End_ and DNA_O-Dyad_), and assembled nucleosomes with these DNA templates (Nuc_O-End_ and Nuc_O-Dyad_; Figure 3A). We then conducted single-molecule TIRF experiments to measure the interaction between Oct4 and these substrates (Figure 3B). The residence time distribution of Oct4 on each substrate can be well described by single-exponential kinetics (Figure 3C). Interestingly, unlike Sox2, the lifetimes of Oct4 on these substrates are statistically identical among each other (Figure 3D and Table S1), suggesting that Oct4 displays no discrimination between bare DNA and nucleosome substrates, or between end-positioned and dyad-positioned nucleosomal motifs (Figures S4D-S4E). We note that in all of the Sox2/Oct4 binding experiments described above, the density of fluorescent nucleosomes on the surface did not decrease after addition of the TF (Figure S5), suggesting that Sox2/Oct4 binding did not eject histones from the DNA.

**Figure 3.**
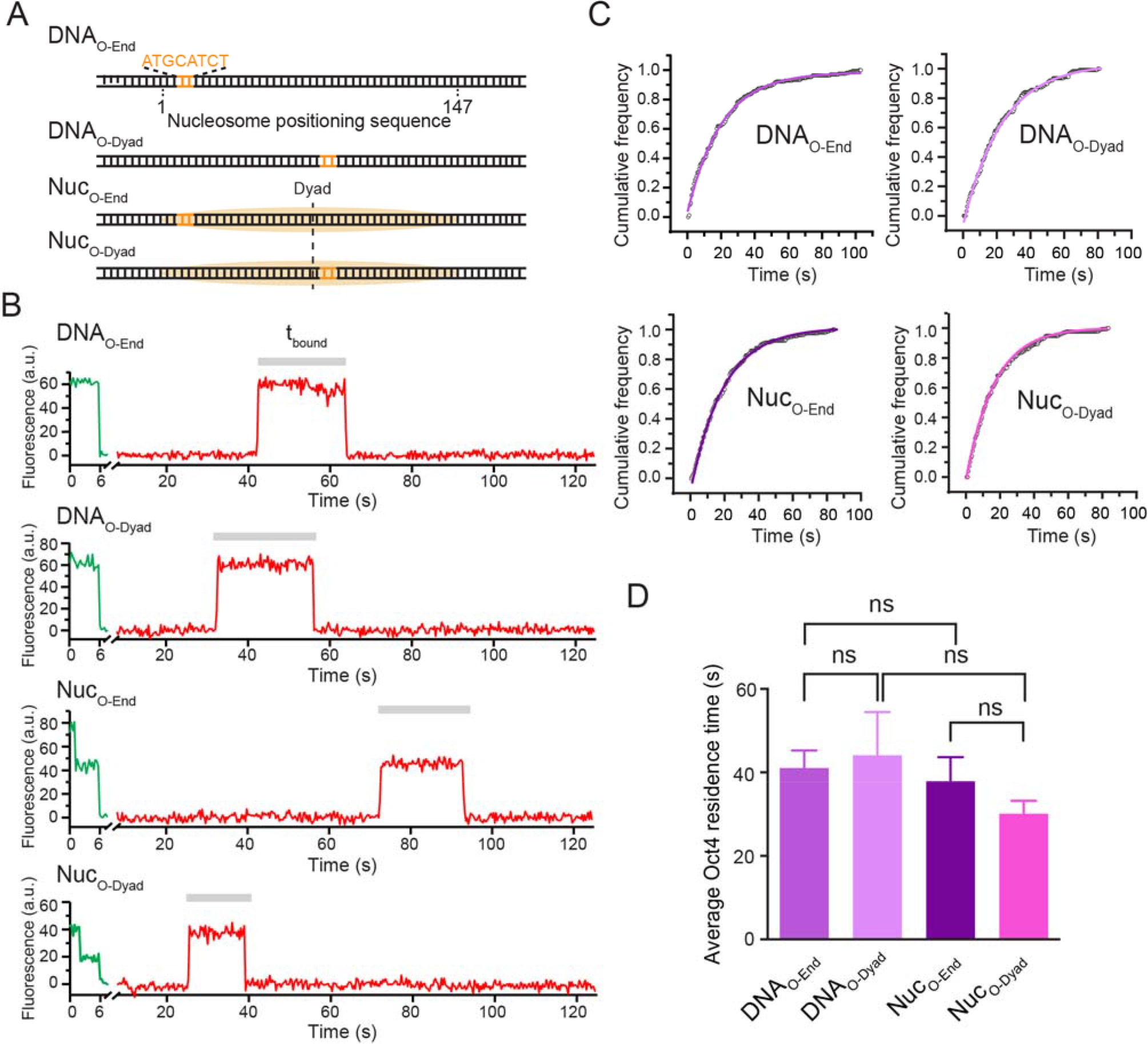
Oct4’s Nucleosome Targeting Activity Is Insensitive to Its Motif Position. (**A**) Diagrams of DNA and nucleosome substrates containing an Oct4 binding motif located at the end or dyad of the 601 nucleosome positioning sequence. (**B**) Representative single-molecule fluorescence trajectories showing Cy5-labeled Oct4 binding to different DNA and nucleosome substrates. (**C**) Cumulative distributions of the Oct4 residence time on different substrates and their respective single-exponential fit. (**D**) A comparison of the average Oct4 residence time on different substrates. Data are represented as mean ± SD.

### Nonreciprocal Regulation Between Sox2 and Oct4 in Nucleosome Binding

Having characterized the individual behaviors of Sox2 and Oct4, next we set out to interrogate the cooperativity between this TF pair in nucleosome targeting. First we engineered a composite Sox2:Oct4 binding motif into the 601 NPS at the end position (nucleotide #1-15) and assembled the nucleosome substrate termed Nuc_SO-End_ (Figure 4A). We then examined the effect of Oct4 on Sox2 binding by complementing Cy5-Sox2 with unlabeled Oct4 in the single-molecule experiments. We found that Oct4 dramatically prolongs the average residence time of Sox2 on Nuc_SO-End_ (Figure 4B). Notably, the stimulatory effect of Oct4 on Sox2 is restricted to the nucleosome substrate, as Oct4 has no effect on Sox binding to the bare DNA substrate DNA_SO-End_ (Figure 4B). A detailed kinetic analysis revealed that the lengthened dwell time of Sox2 on Nuc_SO-End_ by the presence of Oct4 is mainly attributed to an increased weight of the specific mode of Sox2 binding events (A_2_ = 13% without Oct4 vs. 36% with Oct4; Figures 4C-4D; Table S1). Therefore, Oct4 appears to enhance the affinity of Sox2 to an end-positioned nucleosomal target. This result was corroborated by the bulk electrophoretic mobility shift assay (EMSA), which showed that the dissociation constant (*K*_D_) for Sox2-Nuc_SO-End_ interaction is smaller in the presence of Oct4 than in its absence (Figures 4E-4F, S6A).

**Figure 4.**
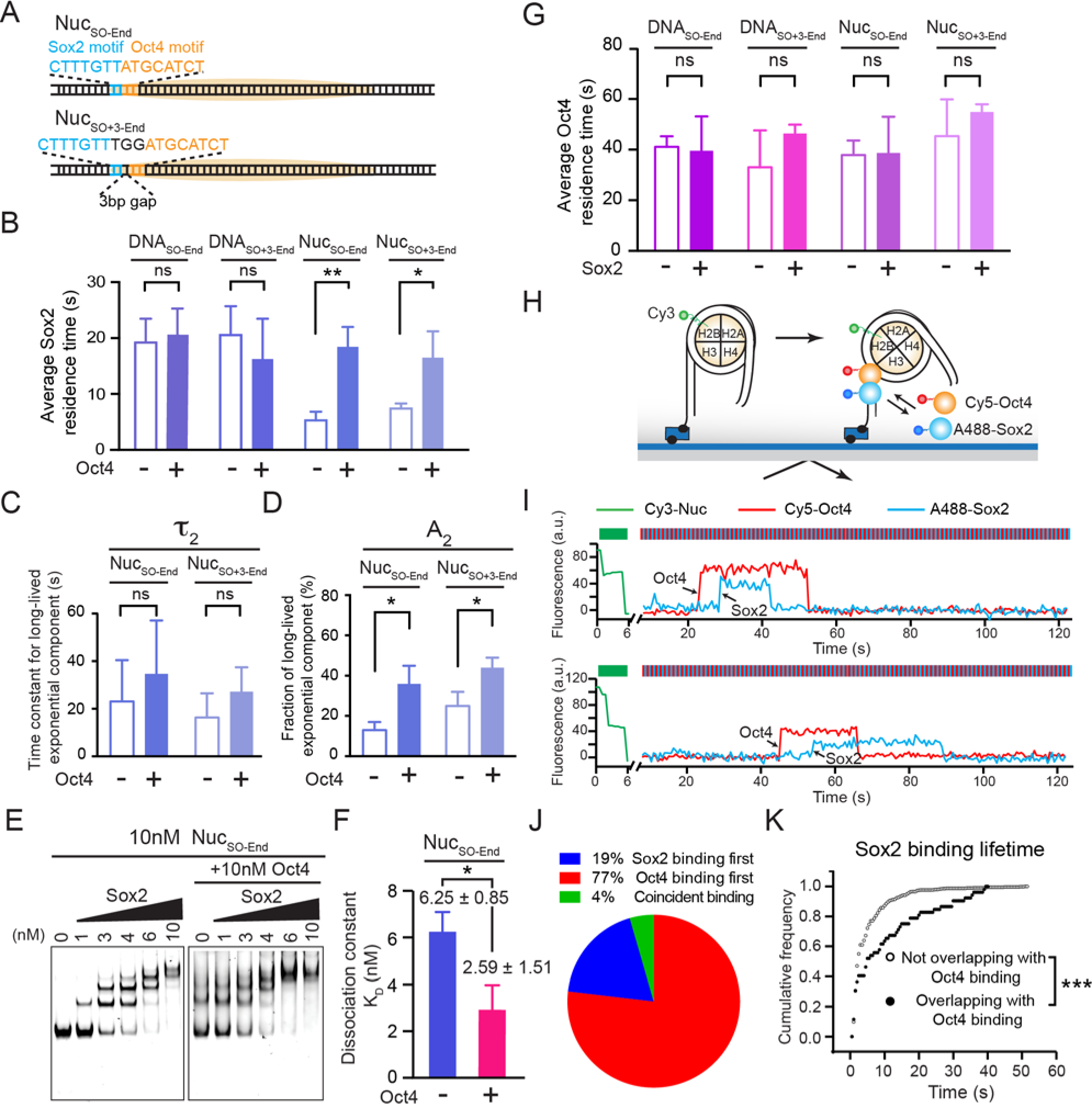
Oct4 and Sox2 Display Nonreciprocal Cooperativity and Hierarchical Engagement in Nucleosome Targeting. (**A**) Diagrams of nucleosome substrates containing an end-positioned Sox2:Oct4 composite motif, either with no gap or with a 3-bp gap between Sox2 and Oct4 binding sites. (**B**) Average residence times of Sox2 on different DNA and nucleosome substrates containing a composite motif in the absence and presence of Oct4. (**C**) Time constants for the long-lived, specific Sox2 binding mode (*τ*_2_) on nucleosome substrates with a non-gapped or gapped composite motif in the absence and presence of Oct4. (**D**) Relative populations of specific Sox2 binding events (A_2_) for different nucleosome substrates in the absence and presence of Oct4. (**E**) A representative EMSA gel showing the formation of Sox2:Nuc_SO-End_ complexes, or the formation of Sox2:Oct4:Nuc_SO-End_ ternary complexes when Oct4 is present, at different Sox2 concentrations. (**F**) Dissociation constants (*K*_D_) for the Sox2:Nuc_SO-End_ interaction in the absence and presence of Oct4 determined from the EMSA results. (**G**) Average dwell times for Oct4 binding to different DNA and nucleosome substrates that contain a composite motif in the absence and presence of Sox2. (**H**) Schematic of the three-color TIRF assay that simultaneously monitors Sox2 and Oct4 binding. Histone H2B, Oct4 and Sox2 are labeled with Cy3, Cy5 and AlexaFluor488, respectively. (**I**) Representative fluorescence-time trajectories showing overlapping Sox2 and Oct4 binding events on Nuc_SO-End_, which reveal the order of TF engagement. (**J**) Pie chart showing the distribution of different scenarios regarding the order of Sox2/Oct4 targeting to nucleosome substrates. (**K**) Cumulative distributions of the lifetime of Sox2 binding events that overlapped with an Oct4 binding event (filled circles) and those that did not overlap (open circles) (*P* = 3.3 × 10^−11^, two-sided Kolmogorov-Smirnov test). Data are represented as mean ± SD.

Besides the canonical composite motif in which Sox2 and Oct4 binding sites are immediately juxtaposed with each other, a variant motif composed of Sox2 and Oct4 sites separated by 3 bp is also found in some cis-regulatory elements such as the *Fgf4* enhancer (Ambrosetti et al., 1997). We generated a nucleosome substrate that contains such a gapped composite motif at its end position (Nuc_SO+3-End_; Figure 4A) and found that Oct4 exerts a similar, albeit somewhat weaker, positive effect on Sox2 engagement (compare Nuc_SO+3-End_ and Nuc_SO-End_, Figures 4B-4D).

We then conducted the converse experiments by using Cy5-Oct4 and unlabeled Sox2 to check the influence of Sox2 on Oct4’s binding activity. Neither DNA nor nucleosome targeting by Oct4 was significantly affected by Sox2 (Figure 4G). Therefore, despite both being classified as PFs, Oct4 can help Sox2 access nucleosomal DNA ends but not vice versa. In other words, the regulation of Sox2 binding by Oct4 is not reciprocal.

### Hierarchically Ordered Targeting of Oct4 and Sox2 to Nucleosomes

To directly follow the order of binding by Sox2 and Oct4 to the same nucleosome target, we labeled the TF pair with distinct fluorophores—Oct4 with Cy5 and Sox2 with AlexaFluor488—and used an alternating laser excitation scheme to simultaneously monitor their behaviors (Figure 4H). We found that the vast majority of overlapping Sox2/Oct4 binding events feature Oct4 binding first followed by Sox2 arrival (Figures 4I-4J). This result, together with the single-color residence time measurements described above, suggests that Oct4 behaves as a pioneer factor that recruits Sox2 and stabilizes its interaction with target sites located at the nucleosome end. Consistently, in the dual-color experiment, the Sox2 binding events that overlapped with an Oct4 binding event are significantly longer than those that did not overlap (Figure 4K).

### Effect of Oct4 on Sox2’s Nucleosome Binding Activity Is Position-Dependent

We then asked whether the enhanced binding of Sox2 in the presence of Oct4 could be observed at other nucleosomal positions besides the end. In particular, we were curious as to whether binding to the nucleosome dyad, which is already preferred by Sox2, could be further stimulated by Oct4. To answer this question, we placed a canonical Sox2:Oct4 composite motif at the dyad position (nucleotide #72-86) of the 601 NPS (Nuc_SO-Dyad_; Figure 5A). Surprisingly, this construct produced a markedly different picture than Nuc_SO-End_: Oct4 has a negligible impact on the average residence time of Sox2 on Nuc_SO-Dyad_ (Figure 5B); it moderately reduces the lifetime of specific Sox2 binding (Figure 5C), but does not affect its relative population (Figure 5D). Moreover, EMSA results showed that *K*_D_ for the Sox2-Nuc_SO-Dyad_ interaction was increased by the presence of Oct4 (Figures 5E-5F, S6B). These results suggest that, instead of promoting Sox2 binding as observed at end-positioned composite motifs, Oct4 weakly diminishes the affinity of Sox2 to the nucleosome dyad. We speculated that such inhibitory effect might be caused by the geometrical interference between the two TFs. Indeed, when a 3-bp gap was inserted between Sox2 and Oct4 binding sites at the dyad position (Nuc_SO+3-Dyad_; Figure 5A), the negative effect of Oct4 on the specific Sox2 binding mode was attenuated (Figure 5C). On the other hand, Sox2 exerted minimal influence on Oct4 binding to dyad-positioned composite motifs (Figure 5G), similar to the results obtained with end-positioned motifs (Figure 4G).

**Figure 5.**
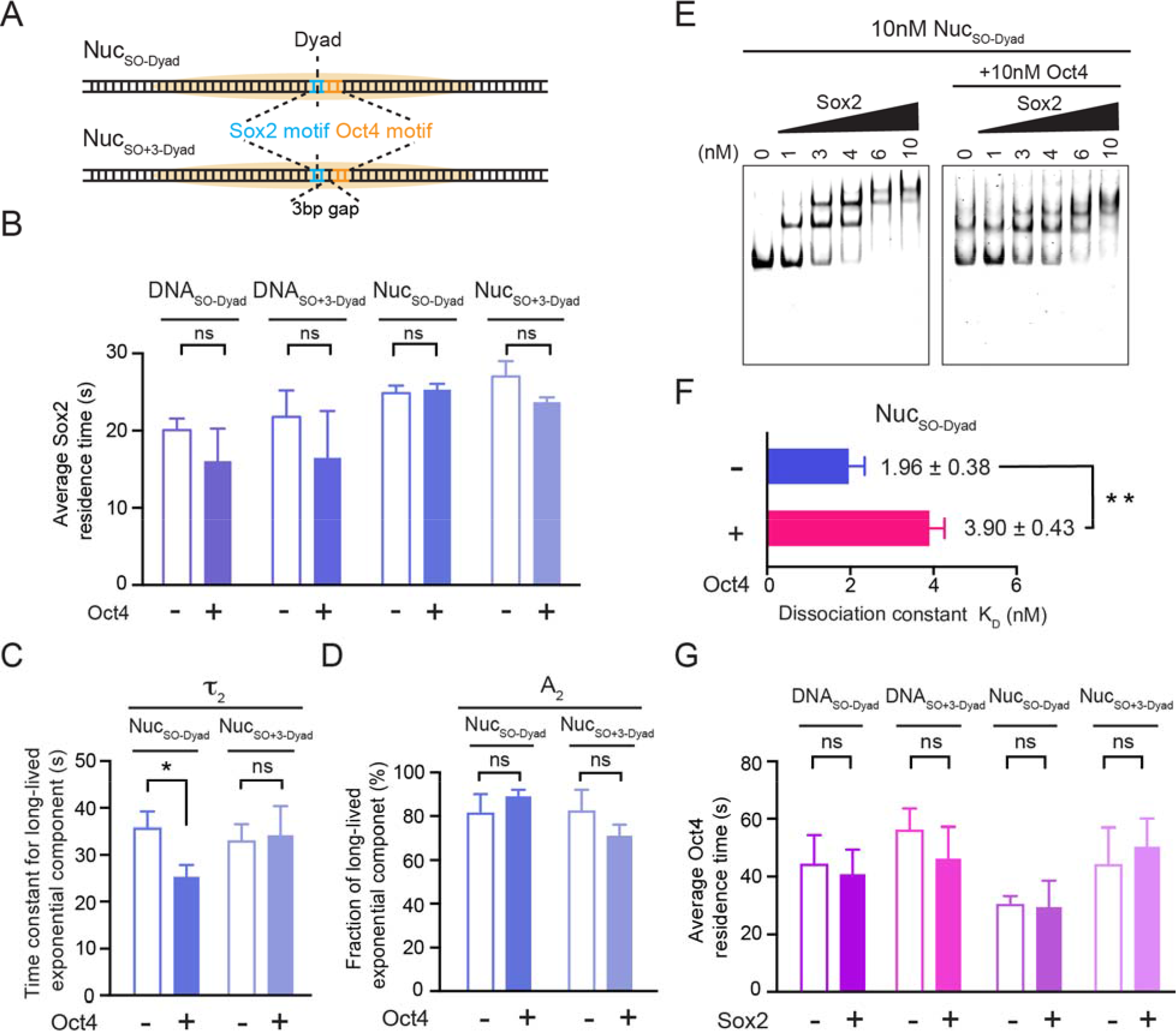
Oct4 Negatively Influences Sox2 Binding to the Nucleosome Dyad. (**A**) Diagrams of nucleosome substrates containing a dyad-positioned Sox2:Oct4 composite motif, either with no gap or with a 3-bp gap between the Sox2 and Oct4 motifs. (**B**) Average residence times of Sox2 on different DNA and nucleosome substrates containing a dyad-positioned composite motif in the absence and presence of Oct4. (**C**) Time constants for the specific Sox2 binding mode (*τ*_2_) on Nuc_SO-Dyad_ and Nuc_SO+3-Dyad_ in the absence and presence of Oct4. (**D**) Relative populations of specific Sox2 binding events (A_2_) for Nuc_SO-Dyad_ and Nuc_SO+3-Dyad_ in the absence and presence of Oct4. (**E**) A representative EMSA gel showing the formation of Sox2:Nuc_SO-Dyad_ complexes at different Sox2 concentrations in the absence and presence of Oct4. (**F**) Dissociation constants (*K*_D_) for the Sox2:Nuc_SO-Dyad_ interaction in the absence and presence of Oct4 determined from the EMSA results. (**G**) Average dwell times for Oct4 binding to different DNA and nucleosome substrates that contain a dyad-positioned composite motif in the absence and presence of Sox2. Data are represented as mean ± SD.

Overall, these results show that the nonreciprocal cooperativity bestowed by Oct4 upon Sox2 could be positive or negative depending on the position and configuration of the composite motif in the nucleosome context.

### Cooperativity between Pluripotency TFs at a Native Genomic Locus

Next we explored the binding behavior of Sox2 and Oct4 at a natural genomic site. We chose the human *LIN28B* locus, which encodes a key protein regulating cell pluripotency and reprogramming (Shyh-Chang and Daley, 2013). We cloned a 162-bp-long DNA segment from this region, which is occupied by a well-positioned nucleosome and targeted by both Sox2 and Oct4 (Soufi et al., 2015). We then used this DNA template to reconstitute nucleosomes, termed Nuc_LIN28B_, and conducted single-molecule binding experiments using fluorescently labeled Sox2 or Oct4. The predicted cognate Sox2 binding site is located between the end and dyad of the LIN28B nucleosome (Figure 6A), as suggested by DNase I footprinting results (Soufi et al., 2015). Accordingly, the average residence time of Sox2 on Nuc_LIN28B_ is longer than that on Nuc_S-End_ but shorter than that on Nuc_S-Dyad_ (Figure 6B). In contrast, Oct4, with its predicted cognate site also located between the end and dyad of the LIN28B nucleosome, exhibited a binding lifetime on Nuc_LIN28B_ indistinguishable from those on Nuc_O-End_ and Nuc_O-Dyad_ (Figure 6C). These results lend further support to our conclusion that Sox2’s pioneer activity is position-dependent, while Oct4’s is not.

**Figure 6.**
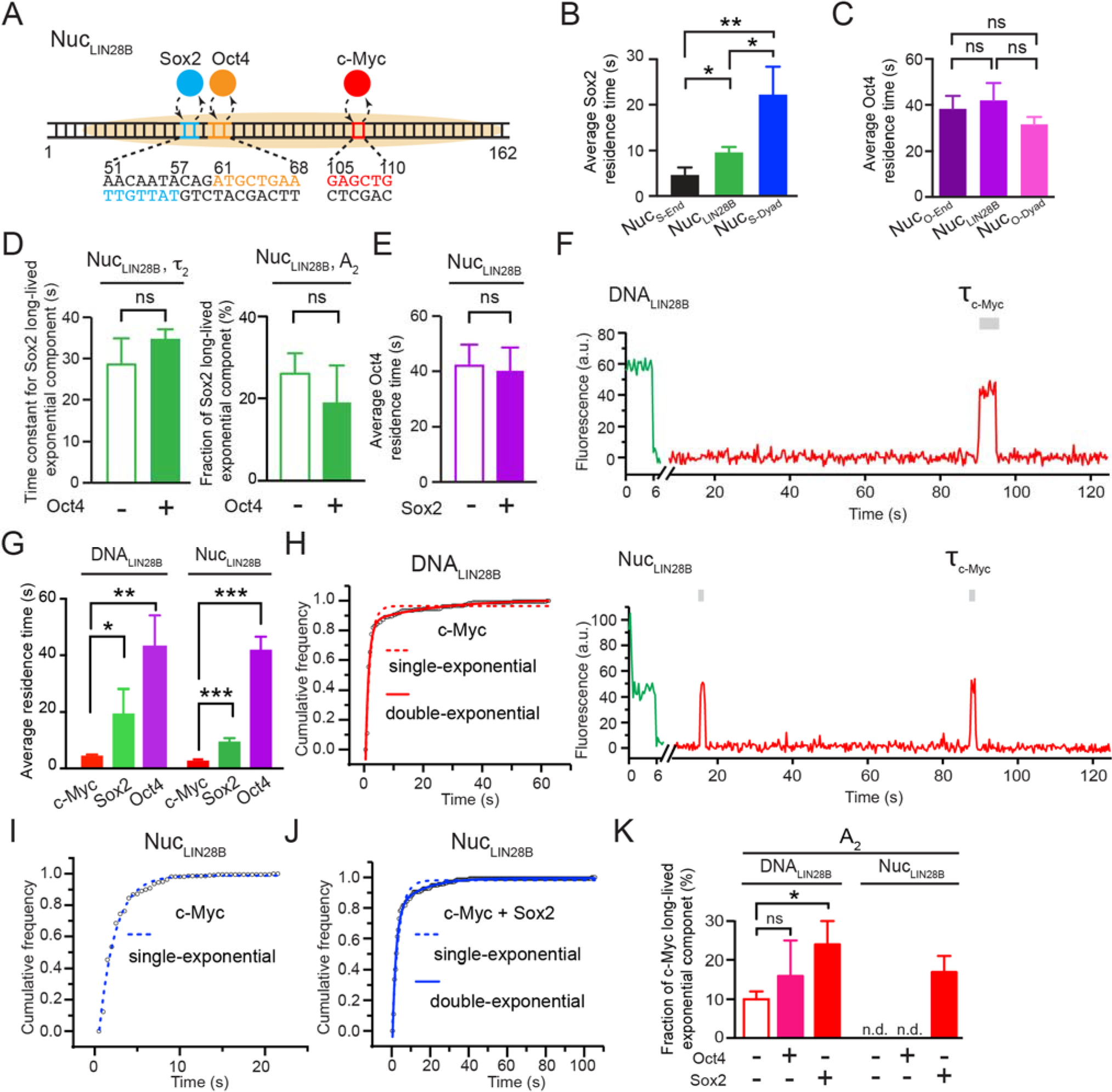
Pluripotency TFs Exhibit Distinct Binding Behaviors at a Native Genomic Locus. (**A**) Diagram of the *LIN28B* genomic locus. Positions of the predicted Sox2, Oct4, and c-Myc binding sites are indicated. (**B**) Comparison of the average residence time of Sox2 on nucleosome substrates with differentially positioned Sox2 binding motifs. (**C**) Comparison of the average residence time of Oct4 on nucleosome substrates with differentially positioned Oct4 binding motifs. (**D**) Time constants (Left) and relative populations (Right) of the specific Sox2 binding mode on Nuc_LIN28B_ in the absence and presence of Oct4. (**E**) Time constants for Oct4 binding to Nuc_LIN28B_ in the absence and presence of Sox2. (**F**) Representative fluorescence-time trajectories showing Cy5-labeled c-Myc:Max binding to Cy3-labeled DNA_LIN28B_ and Nuc_LIN28B_. (**G**) Comparison of the average residence time of each pluripotency TF on DNA_LIN28B_ (Left) and Nuc_LIN28B_ (Right). (**H**) Cumulative distribution (open circles) of the c-Myc residence time on DNA_LIN28B_ and its fit to a single-exponential (dashed curve) or double-exponential function (solid curve). (**I**) Cumulative distribution of the c-Myc residence time on Nuc_LIN28B_ and its fit to a single-exponential function. (**J**) Cumulative distribution of the c-Myc residence time on Nuc_LIN28B_ in the presence of Sox2 and its fit to a single-exponential or double-exponential function. (**K**) Fractions of long-lived c-Myc binding events (A_2_) for c-Myc by itself, in the presence of Oct4, or in the presence Sox2 on DNA_LIN28B_ and Nuc_LIN28B_. Data are represented as mean ± SD.

We then examined Sox2-Oct4 partnership on Nuc_LIN28B_. The Sox2 and Oct4 binding sites within the LIN28B nucleosome are separated by 3 bp and oriented in opposite directions (Figure 6A), unlike all aforementioned composite motifs used in this study that feature a co-directional arrangement. We did not observe significant changes in the Sox2 binding kinetics caused by the addition of Oct4, nor vice versa (Figures 6D-6E). Therefore, besides the position and separation of Sox2/Oct4 binding motifs, their relative direction is also important for cooperativity (Chang et al., 2017).

The *LIN28B* locus is also bound by c-Myc, another member of the Yamanaka factor set that, unlike Oct4 and Sox2, is thought to lack the pioneer activity and preferentially binds to open chromatin by itself (Soufi et al., 2015). We purified and fluorescently labeled c-Myc together with its heterodimeric partner Max (Figure S1), and tested its ability to bind DNA and nucleosome substrates at the single-molecule level (Figure 6F). We found that c-Myc binds to DNA_LIN28B_ much more transiently than Sox2 and Oct4. The same trend was observed on the nucleosome substrate Nuc_LIN28B_ (Figure 6G). Thus, c-Myc possesses an inherent, albeit weak, ability to target nucleosomes. The binding lifetime of c-Myc on DNA and nucleosome substrates is prolonged by the presence of Sox2 and, to a lesser extent, Oct4 (Table S1). Similar to Sox2, c-Myc’s residence time distribution on DNA_LIN28B_ is described by two exponential components (Figure 6H). Interestingly, the long-lived binding component vanished when c-Myc was interacting with Nuc_LIN28B_ (Figure 6I), and was restored by the addition of Sox2 but not Oct4 (Figures 6J-6K). Notably, no Sox2 cognate sequence is found near the specific c-Myc binding site. Hence the stabilizing effect of Sox2 on c-Myc is probably mediated by nonspecific Sox2-nucleosome interaction or by direct Sox2:c-Myc contacts.

### Genome-Wide Binding Preference of Sox2 and Oct4 With Respect to Nucleosome Positioning

The *in vitro* single-molecule data described above revealed the differential positional preference of Sox2 and Oct4 for nucleosomal DNA. To investigate whether such a difference can be recapitulated on a genomic scale, we mined published nucleosome mapping and Sox2/Oct4 ChIP-seq data and interrogated the distributions of Sox2 and Oct4 binding sites relative to nucleosome locations (Figure 7A). We first calculated the aggregate nucleosome-positioning score surrounding Sox2/Oct4 binding sites in mouse embryonic stem cells (mESCs) (Teif et al., 2012) and epiblast stem cells (mEpiSCs) (Matsuda et al., 2017). In both cell types, the average nucleosome occupancy across all Sox2 binding sites is much greater than that for all Oct4 binding sites (Figure 7B), indicating that Sox2 binding sites are more enriched in nucleosome-occupied regions than Oct4 sites.

**Figure 7.**
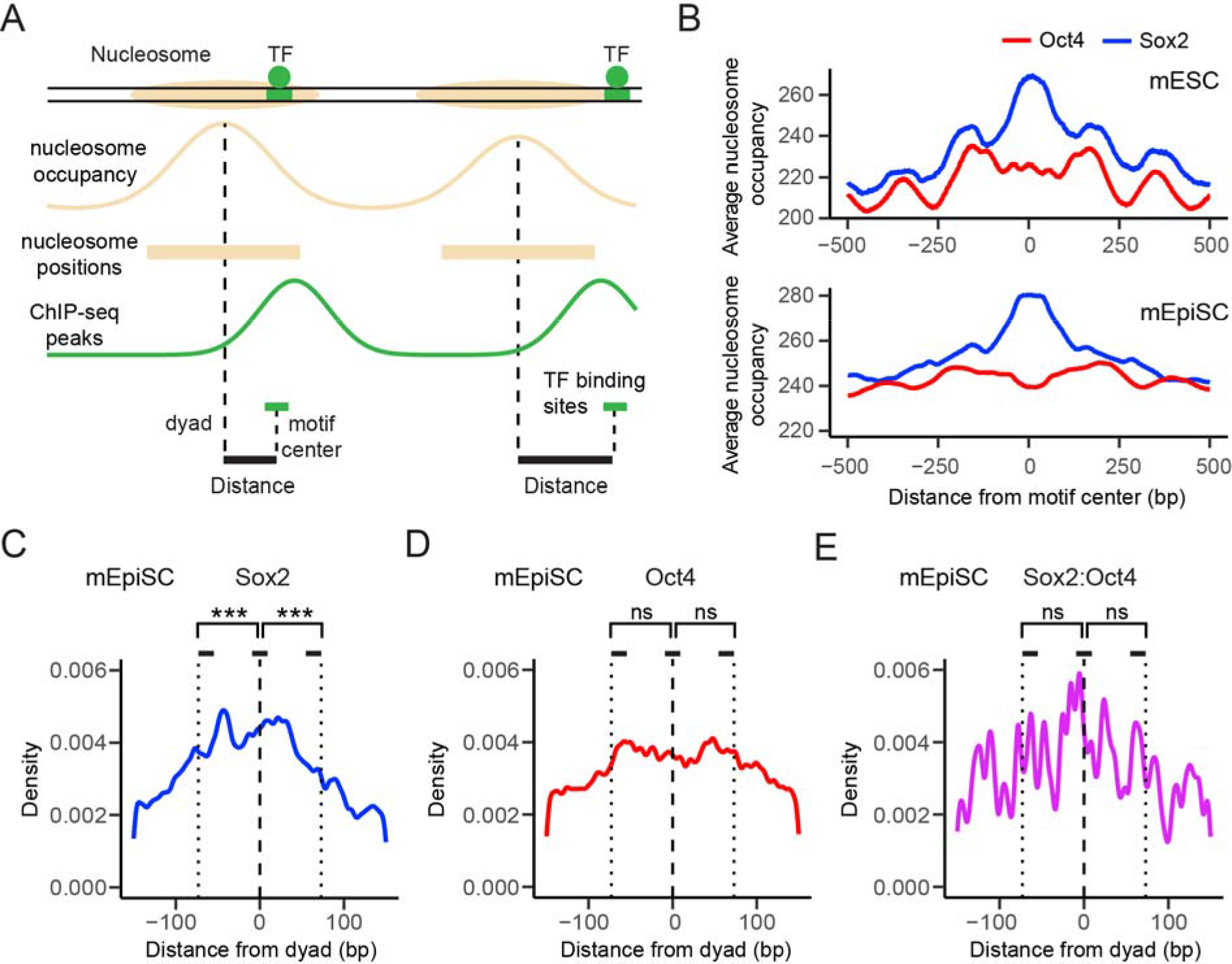
Genome-wide Analyses of the Positional Preference of Sox2 and Oct4 Binding Relative to Nucleosome Positioning. (**A**) Diagram illustrating the procedure of determining genome-wide nucleosome occupancy and TF binding sites. Nucleosome positions were derived from MNase-seq data. TF binding sites were identified by searching for a cognate sequence motif near a ChIP-seq peak for the given TF. (**B**) Nucleosome occupancy scores within a 1,000-bp window surrounding a Sox2 (blue) or Oct4 (red) binding site averaged over all sites identified in mouse embryonic stem cells (top) or epiblast stem cells (bottom). Position 0 corresponds to the center of a given TF binding motif. (**C**) Distribution of the distance between the center of a Sox2 binding site and the nearest nucleosome dyad smoothed with a 3-bp filter. Position 0 (dashed line) corresponds to the dyad; the dotted lines approximate the edges of the nucleosome. *t*-tests were conducted between a 15-bp window centered at the dyad and a 15-bp window inside the nucleosome edge. (**D**) Same as (**C**), except for analyzing the distribution of Oct4 binding sites with respect to a nucleosome. (**E**) Same as (**C**), except for analyzing the distribution of Sox2:Oct4 composite sites with respect to a nucleosome.

We then plotted the distribution of Sox2/Oct4 binding sites in mEpiSCs averaged over all nucleosome-bound regions aligned by their dyad positions. Within the 147-bp window corresponding to the nucleosome, Sox2 binding sites exhibit a strong preference for the dyad region over the edges of the nucleosome (Figure 7C). In contrast, the distribution of Oct4 binding sites within the nucleosome shows no significant difference between nucleosome dyad and ends (Figure 7D). The preference for dyad diminishes in the case of Sox2:Oct4 composite sites as compared to Sox2-alone sites (Figure 7E), suggesting that the strong positional bias of Sox2 binding to nucleosomal DNA is alleviated by the cooperative interaction of Oct4, in line with the single-molecule results. Similar patterns were observed in mouse and human ESCs (Figure S7). Intriguingly, the distributions of noncanonical Sox2 and Oct4 motifs in the human genome, which are enriched in nucleosome-occupied regions as previously reported (Soufi et al., 2015), show distinct patterns from those of canonical motifs. For example, the tendency of dyad targeting vanishes for the noncanonical Sox2 motif that lacks a predominant “G” at the sixth nucleotide position (Figure S7E), which has been suggested to better accommodate nucleosome binding by abolishing DNA distortion (Soufi et al., 2015). These results imply that the genome sequence contains subtle and diverse codes that regulate differential modes of TF targeting.

## DISCUSSION

TF binding positions are usually classified into nucleosome-depleted and nucleosome-enriched regions. In this work, we went beyond such general co-localization analysis by dissecting the dynamical binding pattern of a TF within a nucleosome. Compared to bulk biochemical and genome-wide binding assays, our single-molecule platform affords higher temporal resolution and unique kinetic information, which enabled us to quantitatively determine the cooperativity between pluripotency TFs in the nucleosome context and discover an unexpectedly intricate Sox2-Oct4 partnership. These results are corroborated by analyses of the *in vivo* genomic data, suggesting that the biophysical principles revealed by the *in vitro* reconstituted system may indeed be exploited by the cell to achieve differential gene regulation.

### Distinct Nucleosome Targeting Properties of PFs

TFs generally prefer binding to nucleosome-depleted regions in the genome (Wang et al., 2012), with the exception of PFs that are able to access closed chromatin (Slattery et al., 2014; Zaret and Mango, 2016). The ensuing question then is: does a PF target all nucleosomal DNA sites with equivalent kinetics? We show that, at least for some PFs, the answer is no. In accordance with earlier studies (Liu and Kraus, 2017; Zhu et al., 2018), our results demonstrate that Sox2’s pioneer activity is regulated by both translational and rotational positioning of its cognate motif within the nucleosome. Dyad-positioned sites support prolonged Sox2 binding compared to end-positioned sites. This may be due to the fact that the dyad region, where only one DNA gyre is wound, can better accommodate DNA bending and minor groove widening that are essential for recognition by the HMG domain of Sox2 (Scaffidi and Bianchi, 2001). Moreover, the specific Sox2 binding mode at the dyad region is even more stable than Sox2 interaction with bare DNA (Table S1), indicating additional contacts between the histones and Sox2. As to the rotational setting of the Sox2 binding motif, a more accessible minor groove (facing away from the histone octamer) generally corresponds to a higher affinity (Figures 1I, 2D, S4A-S4C). Similar preference for rotational phasing of binding motifs has been suggested for other TFs such as p53 (Cui and Zhurkin, 2014).

In contrast, we found no such positional preference for Oct4, which indiscriminately targets nucleosomal DNA at different locations. This could be rationalized by the fact that Oct4 contains two DNA-binding domains (POU_S_ and POU_HD_), each recognizing a 4-bp half-motif locating at opposite sides of the DNA helix (Esch et al., 2013). Regardless of the nucleosomal DNA position, one of these half-motifs remains solvent-exposed (Figures S4D-S4E), which is apparently sufficient for stable Oct4-nucleosome interaction. In support of this idea, partial motifs are prevalently found in Oct4 target sites located in nucleosome-enriched genomic regions (Soufi et al., 2015). Therefore, although both regarded as PFs, Sox2 and Oct4 exhibit drastically different nucleosome binding profiles. We note that these interpretations are based on the superposition between the available structures of TF:DNA complexes and unbound nucleosomes. It is also conceivable that the nucleosome structure may be remodeled upon TF engagement.

We found that c-Myc—previously thought not to have an intrinsic nucleosome targeting capability—can nonetheless bind to nucleosomal DNA, albeit transiently. A fully folded bHLHZ domain—the DNA binding domain of c-Myc—occupies a major fraction of the DNA circumference as seen in the c-Myc:DNA structure (Nair and Burley, 2003), incompatible with nucleosome binding. Therefore, the c-Myc binding events observed on nucleosome substrates likely represent a nonspecific mode of interaction that does not require folding of the basic helix 1 in the bHLHZ domain (Soufi et al., 2015), consistent with our kinetic analysis.

With more sensitive methods such as single-molecule imaging being deployed to interrogate TF-chromatin interaction, the list of nucleosome-binding TFs is expected to continue to grow. Our data further suggest that the pioneer activity of these TFs, governed by the structural characteristics of their respective DNA binding domains, is not a binary property but rather falls on a continuous spectrum.

### Nonreciprocal and Conditional TF-TF Cooperativity

Clustered binding of TFs is a hallmark of cis-regulatory elements, such as promoters and enhancers, which integrate multiple TF inputs to direct gene expression. High-throughput methods have been developed to systematically determine the binding patterns of TF pairs on DNA (Chang et al., 2017; Jolma et al., 2015; Siggers et al., 2011; Slattery et al., 2011). However, the biophysical basis for cooperative TF binding in the nucleosome context remains underexplored. In particular, the relationship of chromatin targeting between a PF and a non-pioneer factor, and between a pair of PFs, is still under debate. Using Sox2-Oct4 as a model system, our study sheds new light on this issue. First of all, cooperativity can be unilateral. Oct4 stabilizes the binding of Sox2 to an end-positioned motif, perhaps by opening up a stretch of nucleosomal DNA and generating a local environment permissive to Sox2 binding. Conversely, Sox2 has no effect on Oct4’s behavior on the nucleosome. Using multi-color imaging, we directly followed the order of TF engagement with the nucleosome and found that Oct4 preceding Sox2 is the predominant scenario. This is notably different from live-cell results, which concluded that Sox2 is the lead TF that guides Oct4 to its target sites (Chen et al., 2014). Such discrepancy could be due to the heterogeneous chromatin states inside the cell, which may complicate data interpretation. In any case, the pioneer activity of a TF appears to be hierarchical, reinforcing the aforementioned notion that it should not be considered as an all-or-none trait.

Secondly, we showed that the Sox2-Oct4 cooperativity is strongly dependent on the geometry of the composite motif in the nucleosome context. Contrary to the positive effect exerted at nucleosomal-end positions, Oct4 has a negative impact on Sox2’s access to the dyad region, where Sox2 exhibits a robust pioneer activity by itself. We speculate that, without the benefit of creating extra free DNA surface for Sox2, steric hindrance between the two proteins may become the deciding factor around the dyad region. Therefore, when two TFs are invading the same nucleosome, multiple mechanisms can contribute to their interplay, yielding synergistic or antagonistic binding depending on the specific motif arrangement. The structural determinants of Sox2-Oct4 cooperativity have been studied in depth with DNA substrates (Jauch et al., 2011; Merino et al., 2014; Remenyi et al., 2003; Tapia et al., 2015). Future work tackling the structures of Sox2:Oct4:nucleosome ternary complexes will help illuminate the mechanism by which the nonreciprocal and conditional cooperativity is accomplished.

### Implications for Combinatorial Gene Regulation by TFs

Combinatorial control of gene expression by specific sets of TFs underlies the operation of diverse gene regulatory networks (Thompson et al., 2015). Our single-molecule results, complemented with genomic data analyses, illustrate that the regulatory capacity of TF circuits can be further expanded when integrated into the chromatin context. One could envision that, depending on the location, spacing, and orientation of individual binding motifs within the nucleosome, the same group of TFs can display different nature and strength of cooperativity, thereby activating one set of genes—to varying extents—while at the same time repressing another set of genes. Importantly, TFs need not to directly interact with each other in order to achieve nucleosome-mediated cooperativity, which greatly broadens the potential scope of this mechanism in gene regulation. In this work we used pluripotency TFs as a model system. It will be interesting to examine the differential cooperativity for other TF circuits in diverse cell types.

In conclusion, our study suggests that, besides the chemical space provided by the nucleosome—in the form of post-translational modifications of histones—that is well known to instruct gene expression, the physical organization of chromatin can also be exploited to encode transcriptional logic. Further research along this direction will bring us closer to a complete understanding of how TF-chromatin association and its variation underpins normal cell physiology and disease (Deplancke et al., 2016), and how dynamic and stochastic molecular interactions lead to deterministic and precise gene expression programs.

## ACKNOWLEDGMENTS

We thank Bryan Harada and Rachel Leicher for help with sample preparation, Michael Wasserman for single-molecule data analysis, Xiangwu Ju for the DNase footprinting assay, and other members of the Liu laboratory for discussions. We also thank Abdenour Soufi for sharing the human Sox2/Oct4 ChIP-seq dataset. S.Li was supported by a Tri-Institutional Starr Stem Cell Scholars Fellowship. E.B.Z. was supported by a Medical Scientist Training Program grant from the NIH (T32GM007739) to the Weill Cornell/Rockefeller/Sloan Kettering Tri-Institutional MD-PhD Program. L.Z. was supported by the Robertson Foundation, a Monique Weill-Caulier Career Scientist Award, and an Alfred P. Sloan Research Fellowship (FG-2018-10627). S.Liu was supported by the Robertson Foundation, the Quadrivium Foundation, a Monique Weill-Caulier Career Scientist Award, a March of Dimes Basil O’Connor Starter Scholar Award (#5-FY17-61), a Kimmel Scholar Award, a Sinsheimer Scholar Award, an NIH Pathway to Independence Award (R00GM107365), and an NIH Director’s New Innovator Award (DP2HG010510).

## SUPPLEMENTARY FIGURES AND TABLES

**Figure S1.**
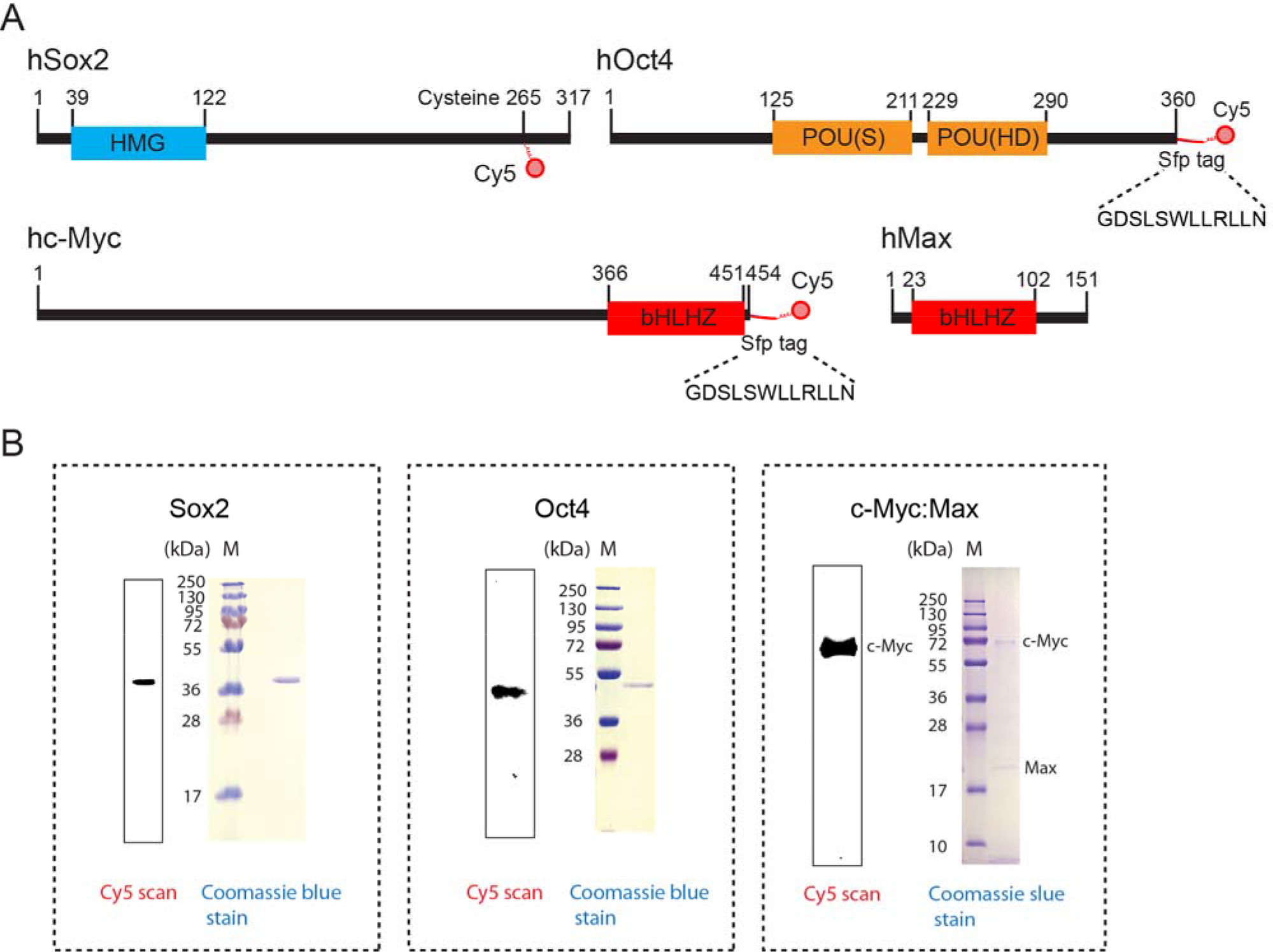
Purification and Site-Specific Labeling of Pluripotency TFs. (**A**) Schematic of the labeling strategies for full-length human Sox2, Oct4, c-Myc, and Max proteins. The DNA-binding domains and their positions are indicated. (**B**) SDS-PAGE analysis of Cy5-labeled Sox2, Oct4, and c-Myc:Max heterodimer.

**Figure S2.**
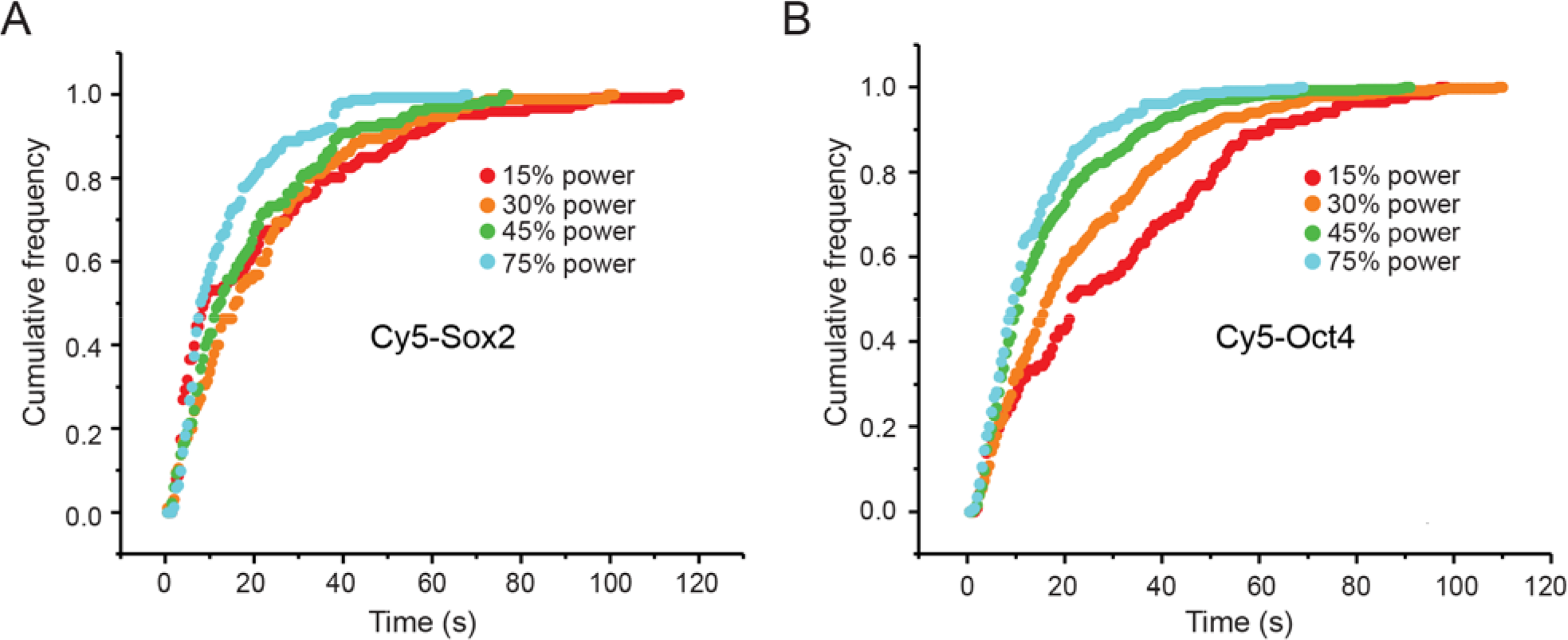
Measuring the Photobleaching Kinetics of Fluorescently Labeled TFs. (**A**) Cumulative distributions of the observed Cy5-labeled Sox2 residence time on DNA measured at different levels of laser power. (**B**) Same analysis for Cy5-labeled Oct4. The photobleaching rate (*k*_bleach_) is assumed to be linearly dependent on the laser power at non-saturating conditions. As such, *k*_bleach_ can be calculated by solving *k*_off,obs_ = *k*_bleach_ + *k*_off_ at multiple laser powers, *k*_off,obs_ and *k*_off_ represent the observed TF dissociation rate constant and true dissociation rate constant, respectively. At 30% power, the time constant for Cy5-Sox2 photobleaching is 75 s; the time constant for Cy5-Oct4 photobleaching is 42 s.

**Figure S3.**
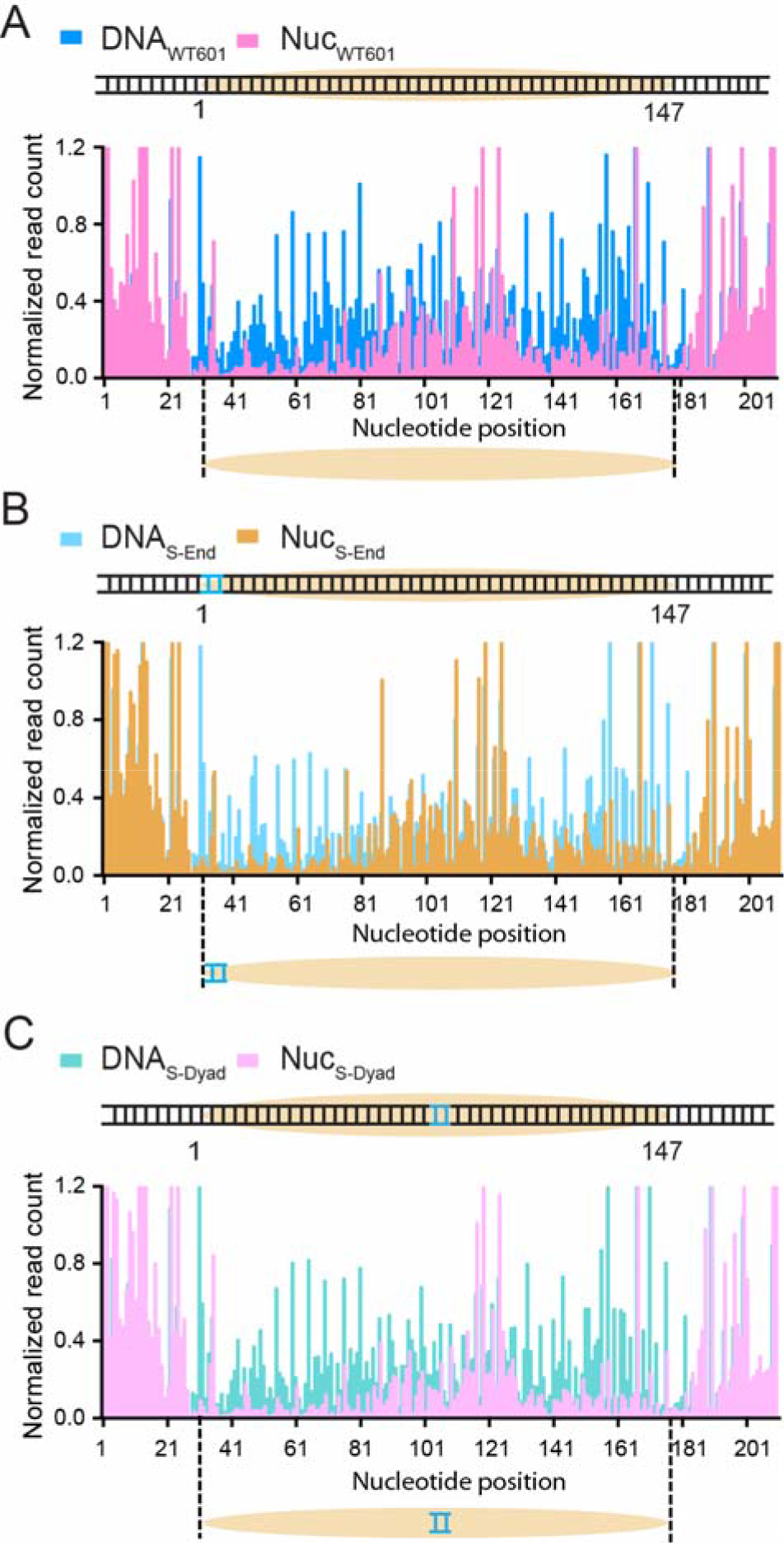
Evaluating Nucleosome Positioning Using a DNase Footprinting Assay. (**A**) DNase I footprinting patterns for a DNA substrate containing a wildtype 601 NPS (DNA_WT601_, blue) and a mononucleosome substrate reconstituted from the same DNA template (Nuc_WT601_, pink). The DNase-protected region in the Nuc_WT601_ data reports the position of the nucleosome. (**B**) Same as (**A**), except with a DNA template containing a 7-bp-long Sox2 binding motif placed at the end of the 601 NPS (DNA_S-End_, light blue; Nuc_S-End_, brown). (**C**) Same as (**A**), except with a DNA template containing a Sox2 binding motif placed at the dyad of the 601 NPS (DNA_S-Dyad_, light green; Nuc_S-Dyad_; light pink). All three nucleosome constructs share an identical protection pattern, suggesting that nucleosome positioning is not perturbed by the engineered DNA sequences.

**Figure S4.**
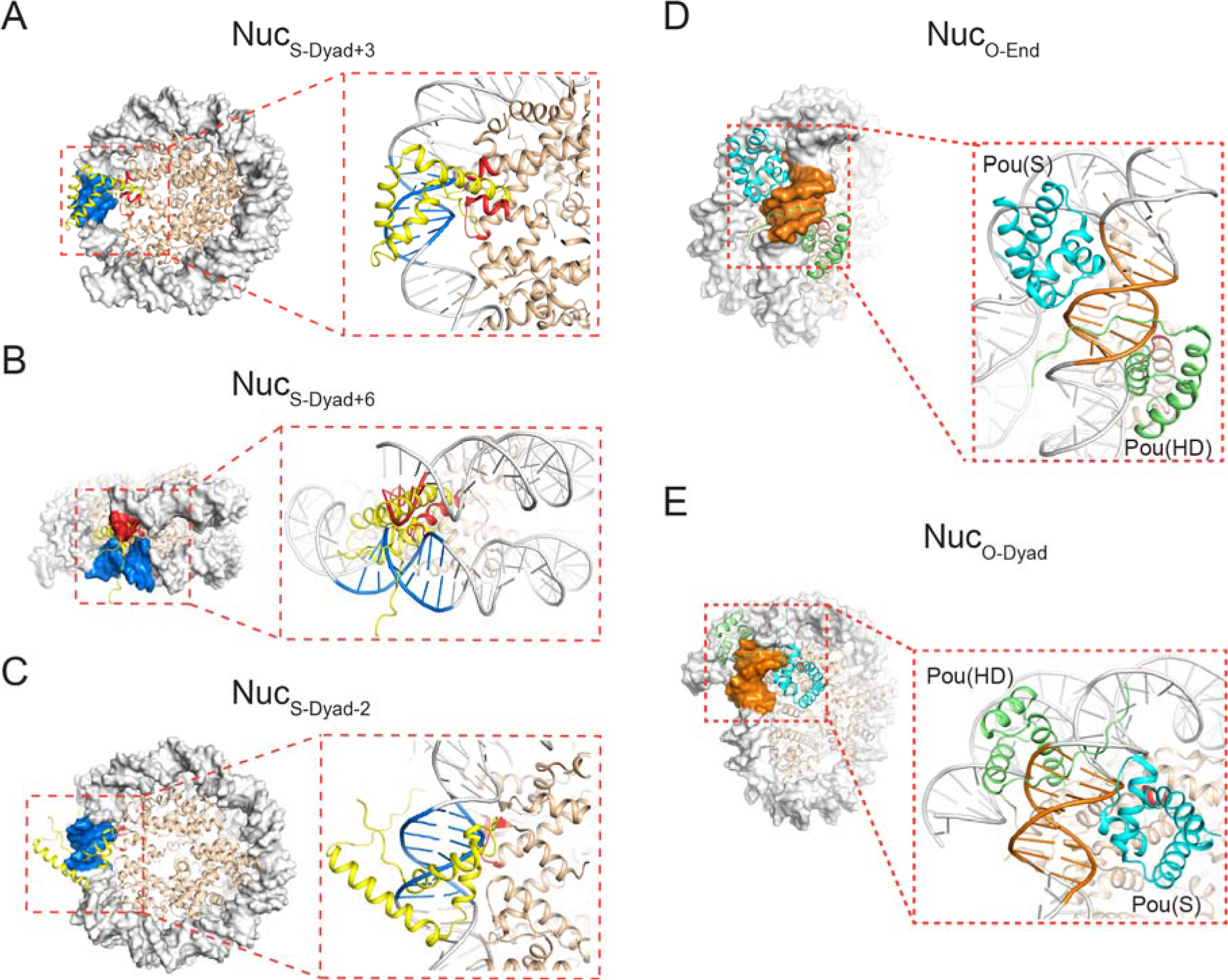
Additional Structural Modeling for Binding Configurations of Sox2 and Oct4 on Nucleosome Substrates. (**A-C**) The Sox2_HMG_:DNA structure (PDB: 1GT0) superimposed on the 601 nucleosome structure (PDB: 3LZ0) aligned by the DNA motif (blue) located at the dyad+3 (**A**), dyad+6 (**B**), and dyad-2 (**C**) positions. Steric clash between Sox2 and the nucleosome is highlighted in red. (**D-E**) Superposition between the Oct4_POU_:DNA structure (PDB: 1GT0) and the 601 nucleosome structure (PDB: 3LZ0) aligned by the DNA motif (orange) located at the end (**D**) and dyad (**E**) positions. The Oct4 POU_HD_ and POU_S_ domains are shown in green and cyan, respectively.

**Figure S5.**
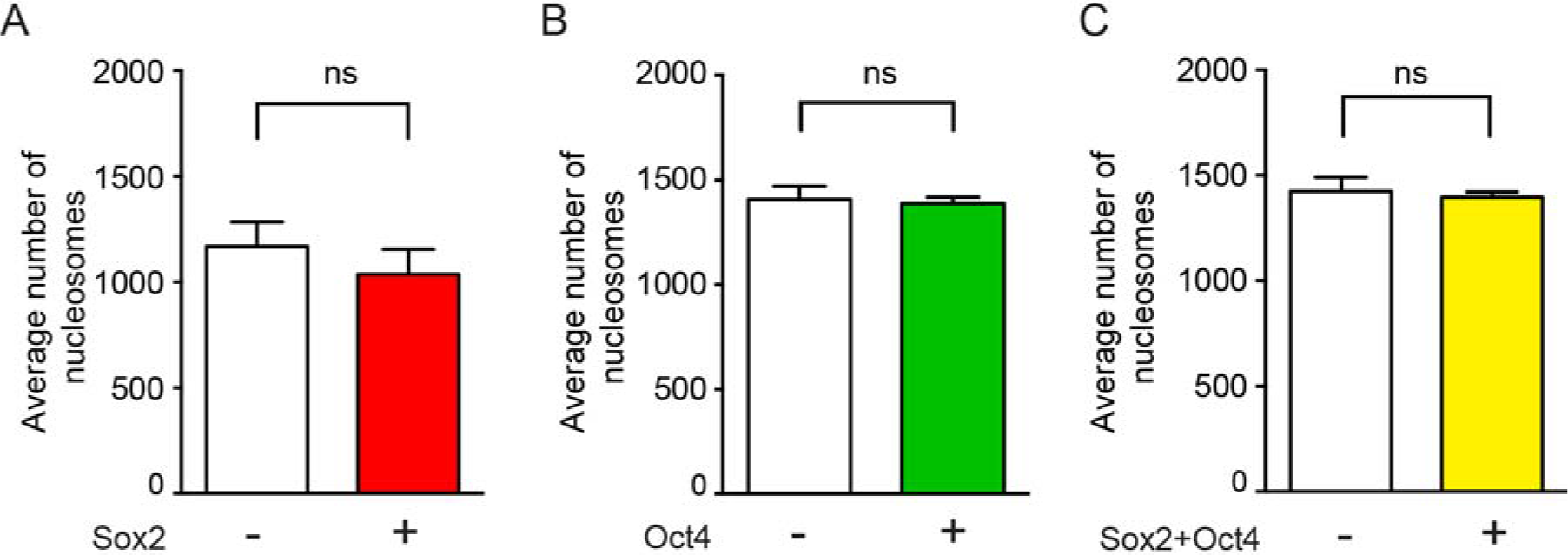
Sox2/Oct4 Binding Does Not Cause Nucleosome Disassembly. (**A**) Average number of surface-immobilized fluorescent nucleosomes containing Cy3-labeled H2B per field of view before (white bar) and 10 minutes after (red bar) the addition of 2 nM Sox2. (**B**) Average number of fluorescent nucleosomes before (white bar) and 10 minutes after (green bar) the addition of 2 nM Oct4. (**C**) Average number of fluorescent nucleosomes before (white bar) and 10 minutes after (yellow bar) the addition of both Sox2 and Oct4. Data are represented as mean ± SD.

**Figure S6.**
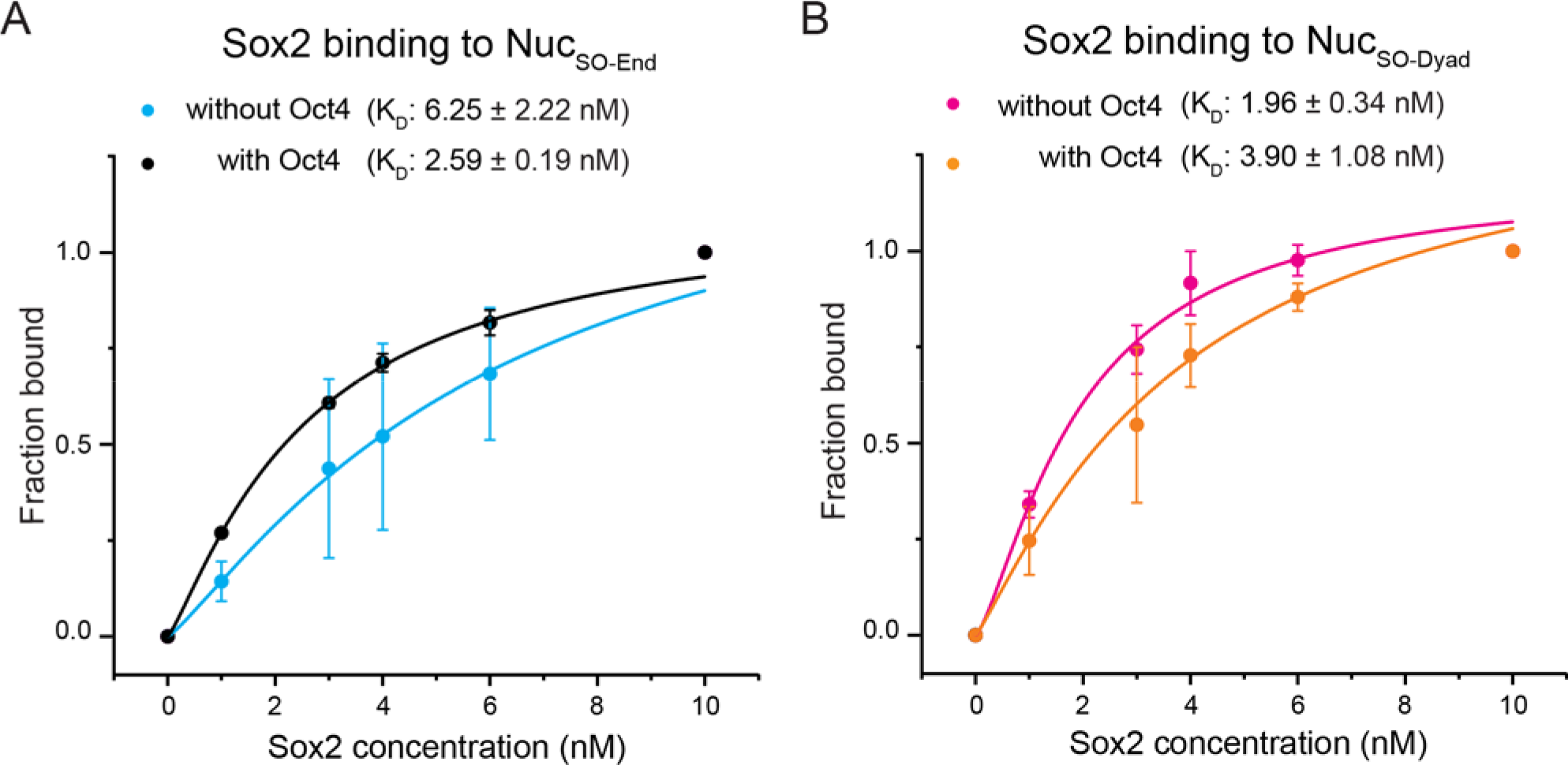
Quantification of Sox2-Nucleosome Interaction from EMSA. (**A**) Fraction of Nuc_SO-End_ substrates that are bound to Sox2 as a function of Sox2 concentration in the absence (blue circles) or presence (black circles) of 10 nM Oct4. *K*_D_ values were determined by fitting the data to a Hill function (blue and black curves). (**B**) Same as (**A**), except for analyzing the Sox2:Nuc_SO-Dyad_ interaction. Data are represented as mean ± SD from three experimental replicates.

**Figure S7.**
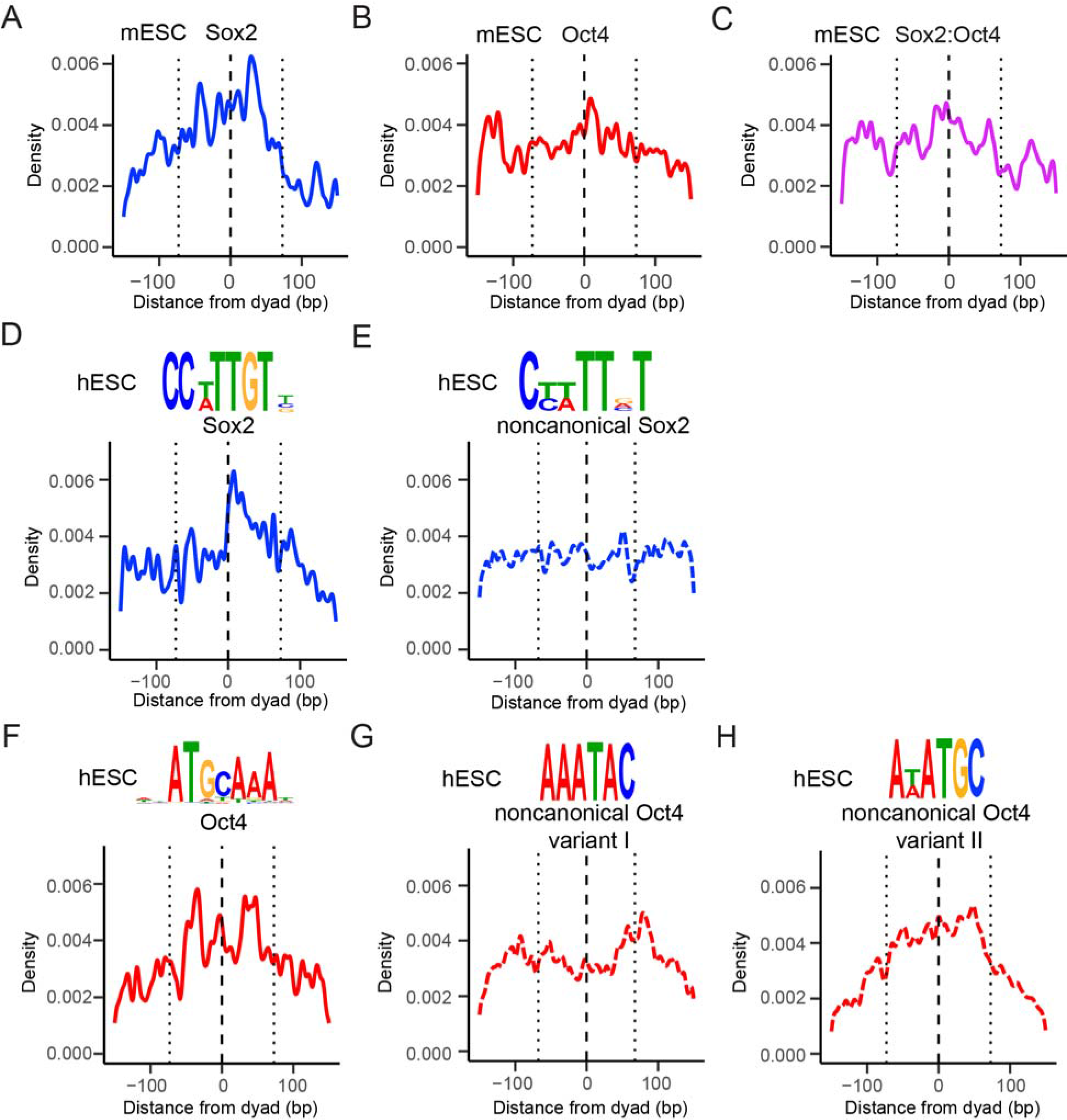
Analyses of Sox2 and Oct4 Binding Preference in Mouse and Human Embryonic Stem Cells. (**A-C**) Distribution of the distance to the nearest nucleosome dyad for Sox2 (**A**), Oct4 (**B**), and Sox2:Oct4 composite binding motifs (**C**) using mESC datasets. Position 0 (dashed line) corresponds to the dyad; the dotted lines approximate the edges of the nucleosome (KS test against uniformity across a 147-bp window corresponding to the nucleosome region, Sox2: *P* < 0.05; Oct4: ns; Sox2:Oct4 composite: ns). (**D-H**) Sequence logo (top) and distribution of the distance to the nucleosome dyad (bottom) for canonical Sox2 binding motifs (**D**), noncanonical Sox2 motifs (**E**), canonical Oct4 binding motifs (**F**), and two variants of noncanonical Oct4 motifs resembling one or the other half of the canonical octameric Oct4 motif (**G** and **H**) using nucleosome-enriched TF binding sites found in hESCs (Soufi et al., 2015). With the exception of the noncanonical Sox2 motif, all motifs feature a non-uniform distribution around the nucleosome region (KS test against uniformity across a 300-bp window centered at the dyad, Sox2: *P* < 0.01; noncanonical Sox2: ns; Oct4: *P* < 10^−4^; noncanonical Oct4 variant I: *P* < 0.01; noncanonical Oct4 variant II: *P* < 2.2 × 10^−16^).

**Table S1.**
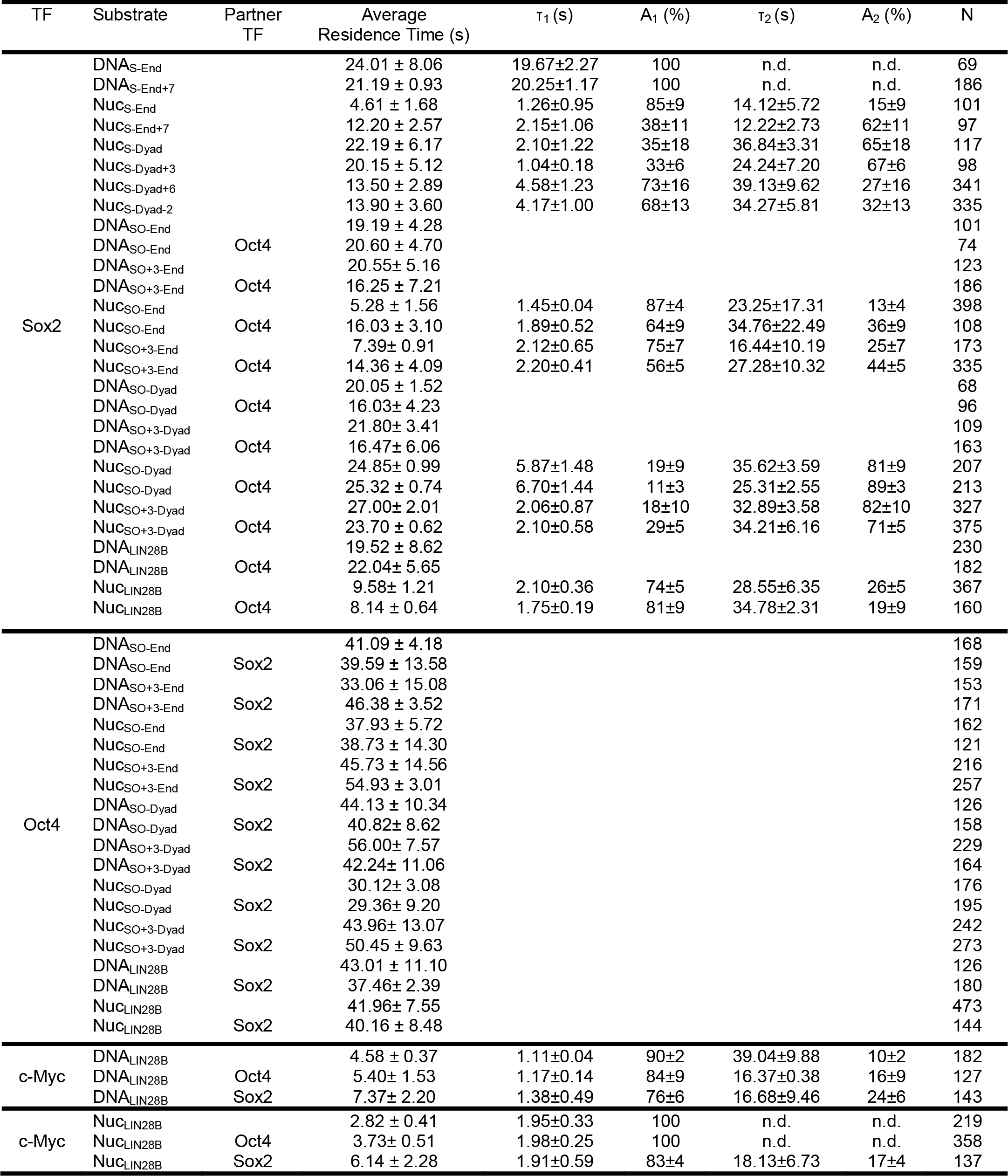
Summary of Kinetic Parameters that Describe TF Binding to Various DNA and Nucleosome Substrates.

## MATERIALS AND METHODS

### Bacterial strains and growth conditions

*E. coli* Rosetta (DE3) plyS cells were cultured in a Luria-Bertani (LB) medium containing 50 μg/ml kanamycin and 34 μg/ml chloramphenicol. *E. coli* BL21 (DE3) cells were cultured at 37 °C in an LB medium containing 100 μg/ml ampicillin.

### Expression, purification, and fluorescent labeling of TFs

The human Sox2 and Oct4 genes were purchased from Addgene and the human c-Myc and Max genes were amplified from HeLa cell cDNA (US Biological T5595-0449). Each gene was cloned into the bacterial vector pET28B with a hexahistidine tag at its N terminus. All proteins were expressed in Rosetta (DE3) plyS cells (Novagen #70956-3) in an LB medium. Cell were grown at 37 °C until OD_600_ reached 0.6, and then induced with 0.5 mM IPTG at 37 °C for 4 hours for Oct4 or 2 hours for Sox2, at 30 °C overnight for c-Myc, or at 25 °C overnight for Max. For Sox2, Oct4 and c-Myc, cells were harvested and lysed by sonication in a denaturing buffer, followed by centrifugation at 15,000 rpm for 40 minutes. The protein was first purified on a Ni-NTA affinity column. The eluted Sox2 and Oct4 were refolded by dialyzing to 2 M urea and then to 0 urea using a desalting column (GE healthcare). Further purification was carried out by gel filtration using a Superdex 200 10/300 GL column (GE Healthcare). Ni-NTA purified c-Myc was mixed with Max at a molar ratio of 1:2.5 as described previously (Farina et al., 2004), and then purified by gel filtration as described above.

Sox2, its only cysteine at amino acid #265 near the flexible C terminus was labeled with Cy5 maleimide (GE healthcare) or AlexaFluor488 C_5_ maleimide (Thermo Fisher) at a mixing ratio of 1:1.5 after desalting column purification. An Sfp tag was introduced to the N terminus of Oct4 and c-Myc (Yin et al., 2005). After desalting column purification, the Sfp-tagged proteins were incubated with the Sfp synthase (purified in-house) and CoA-Cy5 (synthesized and purified in-house) at a molar ratio of 1:1.5:2.5 at 4 °C overnight. To synthesize CoA-Cy5, the coenzyme A trilithium salt (Sigma-Aldrich) was conjugated with Cy5 maleimide at room temperature for 2 hours and purified with a C18 250x 4.6mm column (Agilent) as previously described (Yin et al., 2006). Free dye molecules and the Sfp synthase were removed from the labeled proteins by gel filtration.

### Preparation of histones and DNA templates and nucleosome assembly

*Xenopus laevis* histones were recombinantly expressed in BL21 (DE3) cells. H2B T49C mutant was generated by site-directed mutagenesis. The mutant histone was purified and labeled with Cy3 maleimide (GE Healthcare) under denaturing conditions (Harada et al., 2016). Histone octamers were reconstituted with equal ratio of each histone and purified by gel filtration as described previously (Luger et al., 1999). DNA templates were made by PCR using biotinylated primers and a plasmid containing a 601 NPS (Li et al., 2010; Lowary and Widom, 1998) that was modified such that a Sox2 motif (CTTTGTT), an Oct4 motif (ATGCATCT), a composite Sox2:Oct4 motif (CTTTGTTATGCATCT), or a composite Sox2:Oct4 motif with a 3-bp spacer (CTTTGTTTGGATGCATCT) was placed at indicated positions. The *LIN28B* genomic DNA fragment was synthesized from IDT. The DNA products were purified by ion-exchange chromatography on a Mono Q column (GE Healthcare) and stored in TE buffer (10 mM Tris-HCl, pH 8.0, 0.1 mM EDTA). For fluorescent labeling, DNA modified with a primary amine group was mixed with Cy3 NHS ester (GE Healthcare) at room temperature for 2 hours. Free dyes were subsequently removed by a Sephadex G-25 column (GE Healthcare). Nucleosomes were assembled by salt gradient dialysis as previously described (Lee and Narlikar, 2001). The assembly efficiency was optimized by titrating the DNA:octamer molar ratio. The purity of the products was evaluated on a 5% native TBE-PAGE gel.

### Single-molecule experiments

Glass slides and coverslips were cleaned by sonication in acetone and 1 M KOH, followed by treatment with Nanostrip (VWR). They were then subjected to argon plasma cleaning (Harrick Plasma) followed by silanization with 2% 3-aminopropyltriethoxysilane in acetone. Cleaned slides and coverslips were passivated with a mixture of polyethylene glycol (PEG) and biotin-PEG (Laysan Bio) for 2 hours, followed by a second round of PEGylation with 4 mM MS4-PEG (Thermo Fisher Scientific) for 1 hour. The assembled flow chamber was infused with 40 μl of 0.2 mg/ml streptavidin (Thermo Fisher Scientific), incubated for 5 minutes, and washed with 100 μl of T50 buffer (10 mM Tris-HCl, pH 7.5, 50 mM NaCl). Biotinylated DNA or nucleosome molecules were injected into the chamber and immobilized through streptavidin-biotin linkage. Single-molecule imaging was conducted on a total-internal-reflection fluorescence microscope (Olympus IX83 cellTIRF) equipped with an EMCCD camera (Andor iXon Ultra897). The imaging buffer contained 40 mM Tris-HCl, 12 mM HEPES, pH 7.5, 60 mM KCl, 3 mM MgCl_2_, 10% (v/v) glycerol, 0.02% (v/v) Igepal CA-630, 0.1 mg/ml BSA, and an oxygen scavenging system [1% (w/v) glucose, 1 mg/ml glucose oxidase, 0.04 mg/ml catalase, 2 mM trolox (Sigma-Aldrich)]. Movies were recorded at room temperature with a frame rate of 300 ms. Positions of the immobilized substrates were determined during the initial 20 frames using a 532-nm laser. TF binding dynamics were then observed using a 640-nm laser. To simultaneously monitor two fluorescently labeled TF species, an alternating excitation scheme was employed in which a 640-nm laser and a 488-nm laser were each switched on for one frame in an interlaced fashion. Labeled and unlabeled TFs were typically used at a concentration of 2 nM and 20 nM, respectively. Single-molecule fluorescence-time trajectories were extracted and analyzed by the SPARTAN software (Juette et al., 2016). TF binding events were identified using a fluorescence intensity threshold. Histogram building and curve fitting were performed with the Origin software (OriginLab).

### Electrophoretic mobility shift assay (EMSA)

10 nM of DNA or nucleosome substrates were incubated with indicated TFs in a binding buffer (10 mM Tris-HCl, pH 7.5, 1 mM MgCl_2_, 1 mM DTT, 10 mM KCl, 0.5 mg/ml BSA, and 5% glycerol) at room temperature for 10 minutes. The reaction mixture was loaded on a 5% non-denaturing polyacrylamide gel, which was run in 0.5× Tris-Borate-EDTA at 4 °C at 80 volts, stained by SYBR Gold (Invitrogen), and scanned by a Typhoon FLA 7000 gel imager (GE Healthcare). Band intensities were extracted by ImageQuant (GE healthcare). The fraction of substrates bound by Sox2 is defined as: *I*_Sox2:substrate_/(*I*_Sox2:substrate_ + *I*_free substrate_). The fraction of substrates bound by Sox2 in the presence of Oct4 is defined as: (*I*_TF:substrate_ − *I*_Oct4:substrate_)/(*I*_TF:substrate_ + *I*_free substrate_). *I*_TF:substrate_ denotes the intensity for substrates bound by any TF; *I*_Oct4:substrate_ denotes the intensity for substrates bound by Oct4 alone.

### DNase footprinting

200 ng of bare DNA or reconstituted nucleosome substrates were treated with 0.008 units of DNase I (Invitrogen) at 25 °C for 3 minutes in 50 μl of Buffer I (10 mM Tris-HCl, pH 7.5, 1 mM MgCl_2_, 10 μM ZnCl_2_, 0.2 mM DTT, 10 mM KCl, 0.5 mg/ml BSA, and 5% glycerol) and 50 μl of Buffer II (10 mM MgCl_2_ and 5 mM CaCl_2_). Then the reaction was stopped by the addition of 90 μl of Buffer III (20 mM Tris-HCl, pH 7.5, 50 mM EDTA, 2% SDS, and 0.2 mg/ml proteinase K) and chilled on ice bath for 10 minutes. The digested DNA was cleaned by TE-buffer-saturated phenol:chloroform:isoamyl alcohol (25:24:1, v/v) and then prepared for Illumina sequencing with the NEBNext Ultra II DNA Library Prep Kit (New England BioLabs). The adaptor-ligated DNA was amplified for 10 cycles following the manufacturer’s protocol. Sequencing was performed on a MiSeq platform. The paired-end reads were aligned to the correspondent DNA sequence. The read counts at each nucleotide position across the template sequence were extracted with custom Perl scripts.

### Structure alignment

The structures of DNA in complex with Oct4_POU_ and Sox2_HMG_ (PDB: 1GT0) and the 601-nucleosome (PDB: 3LZ0) were obtained from the RCSB protein data bank (Remenyi et al., 2003; Vasudevan et al., 2010). The DNA bound to Sox2_HMG_ or Oct4_POU_ was superimposed on specific positions of the nucleosomal DNA using the “align” command in PyMOL for aligning the 3’-, 4’-, and 5’-carbon atoms in the ribose sugar.

### Nucleosome positioning

Raw sequencing paired-end reads were downloaded from the NCBI Sequence Read Archive (SRA) for MNase-seq nucleosome mapping data from human ESCs (SRP028172) (Yazdi et al., 2015) and MNase-seq data from mouse ESCs (SRX187610) (Teif et al., 2012). Using bowtie1 (v. 1.2.2) (Langmead et al., 2009) in “-v” alignment mode, raw sequencing reads with a maximum of 2 mismatches were aligned to the appropriate reference assembly (GRCh38 for human data, and mm9 for mouse data), excluding nonstandard chromosomes. Only reads mapping to unique locations were kept for further analysis. DANPOS2 in “dpos” mode (v. 2.2.2) (Chen et al., 2013) was used to infer nucleosome positioning from mapped paired-end reads pooled across runs. Positions were represented as both genomic intervals defining peak locations as well as quantitative nucleosome position scores for 10 bp non-overlapping windows along the genome. Peaks were filtered using a custom script (https://github.com/LiZhaoLab/Oct4Sox2_nuclpos/tree/master/scripts/danpos_xls_process.py) to retain only those with a summit height of 1.5 times the genomic mean. Nucleosome dyads are assumed to correspond to the summit positions.

### TF binding sites

For mouse ESCs, ChIP-seq data were downloaded as raw single-end reads from the NCBI SRA (Oct4, SRR713340; Sox2, SRR713341; control, SRR713343) (Whyte et al., 2013). Fastq data were aligned using hisat2 (v. 2.0.5) (Kim et al., 2015) with default parameters, except with splice-awareness disabled, to the UCSC mm9 assembly of the mouse genome, excluding unmapped and random chromosomes. ChIP-seq peak locations were then called using MACS2 (v. 2.1.2) (Zhang et al., 2008) using the same settings as previously described (defaults except genome size 1.87×10^9^, *P*-value threshold 10^−9^, and “--keep-dup” set to “auto”) (Whyte et al., 2013). For mouse EpiSCs, ChIP-seq data were downloaded as BED intervals from the NCBI GEO (Oct4, GSM1924747; Sox2, GSM1924746) (Matsuda et al., 2017). We defined TF binding sites as locations conforming to the canonical binding motif within 200 bp of the respective ChIP-seq peak; tandem Sox2:Oct4 binding sites were defined as locations conforming to the composite motif within a 400-bp window centered on the overlap of the two ChIP-seq peaks. Genomic sequences around the ChIP-seq peaks from each cell type were then checked for TF binding motifs using FIMO (v. 4.11.2) (Grant et al., 2011) with default parameters and motifs from JASPAR (Oct4, MA1115.1; Sox2, MA0143.3; Sox2:Oct4 tandem, MA0142.1).

For human ESCs, the locations of ChIP-seq peaks found to be nucleosome-enriched (Soufi et al., 2015) were converted from hg19 to GRCh38 via UCSC liftOver (http://hgdownload.cse.ucsc.edu/downloads.html) (Hinrichs et al., 2006). The same 200-bp peak-centered windows as previously used (Soufi et al., 2015) were then checked for canonical TF binding motifs using FIMO as described above (JASPAR motifs: Oct4, MA1115.1; Sox2, MA0143.3). To identify noncanonical binding sites, these same windows were also checked for the noncanonical motifs identified by (Soufi et al., 2015), with a *P*-value threshold of 0.001.

### Statistical analysis

Statistical significance was determined by unpaired two-tailed Student’s *t*-tests using GraphPad Prism. Welch’s two-sample *t*-tests were conducted using the base implementation in the “stats” package of R (v. 3.4.3) on the count values for the central 15 bp and the 15 bp just inside the left and right bounds of the nucleosome (reported *P* values are the greater of the two sides). Kolmogorov-Smirnov tests were performed using the base implementation in the “stats” package of R, except when comparing two different distributions, the bootstrap KS test (“ks.boot”) from the “Matching” package (v. 4.9.3) (Sekhon, 2011) was used instead. The difference between two groups was considered statistically significant when the *P* value is less than 0.05 (\**P* < 0.05; \*\**P* < 0.01; \*\*\**P* < 0.001; ns, not significant).

